# The SON RNA splicing factor is required for intracellular trafficking that promotes centriole assembly

**DOI:** 10.1101/2021.01.29.428802

**Authors:** Alexander J. Stemm-Wolf, Eileen T. O’Toole, Ryan M. Sheridan, Jacob T. Morgan, Chad G. Pearson

## Abstract

Control of centrosome assembly is critical for cell division, intracellular trafficking and cilia. Regulation of centrosome number occurs through the precise duplication of centrioles that reside in centrosomes. Here we explored transcriptional control of centriole assembly and find that the RNA splicing factor SON is specifically required for completing procentriole assembly. Whole genome mRNA sequencing identified genes whose splicing and expression are affected by the reduction of SON, with an enrichment in genes involved in the microtubule cytoskeleton, centrosome and centriolar satellites. SON is required for the proper splicing and expression of *CEP131* which encodes a major centriolar satellite protein and is required to organize the trafficking and microtubule network around the centrosomes. This study highlights the importance of the distinct microtubule trafficking network that is intimately associated with nascent centrioles and is responsible for procentriole development.

## Introduction

Centrosomes are the preeminent nucleators and organizers of microtubules in dividing cells. Within the centrosome is a pair of microtubule-based centrioles that serve both to recruit the surrounding pericentriolar material necessary for microtubule nucleation and to facilitate control of centrosome assembly such that cells have one centrosome at the beginning of the cell cycle and two centrosomes prior to cell division. This promotes the assembly of a bipolar spindle during mitosis, thus ensuring high-fidelity segregation of chromosomes to daughter cells (Loncarek & Bettencourt-Dias, 2018; Nigg & Holland, 2018). Dysregulation of centrosome number can instigate cellular transformation to a cancerous state (Levine et al., 2017). More than two centrosomes results in chromosomal instability (Ganem et al., 2009; Silkworth et al., 2009; Zhang et al., 2015), altered cell migration (Godinho et al., 2014) and disruptions to the microtubule network (Godinho et al., 2014; Marteil et al., 2018; Sankaran et al., 2020). Differentiated cells, on the other hand, often utilize non-centrosomal microtubule organizing centers (MTOCs) and must attenuate the microtubule nucleating function of the centrosome (Sanchez & Feldman, 2017), suggesting the existence of broad programs to regulate MTOC activity. Control of centriole assembly and centrosome activity, therefore, has profound implications for development and disease.

Centriole assembly occurs in distinct stages: assembly is stimulated by the activity of the PLK4 kinase which coordinates the addition of early structural factors that establish the nine-fold symmetry of the structure at its base (Kitagawa et al., 2011; Kleylein-Sohn et al., 2007). Additional factors, such as Centrobin, CPAP and POC5, are required for centriole elongation (Azimzadeh et al., 2009; Gudi et al., 2011; Y.-N. Lin et al., 2013; Schmidt et al., 2009). Protein components required for assembling centrioles traffic in cytoplasmic particles associated both with the centrosome scaffold protein, Pericentrin (PCNT) and with centriolar satellites (Dammermann & Merdes, 2002; Hori et al., 2015; Ito et al., 2019; Kodani et al., 2015; Prosser & Pelletier, 2020). Both Pericentrin granules and centriolar satellites require microtubules and dynein for their accumulation near the centrosome (Kubo et al., 1999; Young et al., 2000). Each stage of the centriole assembly process is a potential target for regulation of nascent centriole assembly.

Centrosome number is controlled by a multi-faceted regulation of centriole duplication coordinated with the cell cycle. This includes PLK1-dependent licensing during mitotic exit that promotes centriole disengagement (Colicino & Hehnly, 2018; Matsuo et al., 2012; Schöckel et al., 2011; Tsou et al., 2009). In S phase, the centriole assembly master regulator, PLK4, initiates early steps in procentriole assembly promoting the removal of the centriole assembly inhibitor Cdc6 (Xu et al., 2017) and focusing centriole assembly to a single site on the mother centriole (Leda et al., 2018; J.-E. Park et al., 2019; Yamamoto & Kitagawa, 2019). Interactions between PLK4 and STIL serve to stimulate PLK4 kinase activity and STIL phosphorylation by PLK4, allowing STIL to interact with the other early centriole assembly factors, SAS-6 and CPAP, initiating the assembly of the early procentriole. (Liu et al., 2018; McLamarrah et al., 2018; Tyler C. Moyer et al., 2015; Tyler Chistopher Moyer & Holland, 2019; Ohta et al., 2018). Furthermore, PLK4 phosphorylates the major centriolar satellite components CEP131 and PCM1, and PLK4-dependent phosphorylation of PCM1 is required for pericentrosomal localization of centriolar satellites (Hori et al., 2016; D. H. Kim et al., 2019), suggesting that trafficking of assembly factors to the centrosome may also be controlled by PLK4 regulation.

Centriole assembly is also regulated at the transcriptional level where distinct mRNA isoforms both promote and repress nascent centriole assembly. Alternative polyadenylation of *CEP135* generates both a short and long isoform of this centriole assembly factor, which have antagonistic effects on centriole assembly (Ganapathi Sankaran et al., 2019). Overexpression of the full-length isoform increases centriole number while overexpression of the short isoform reduces centriole number (Dahl et al., 2015). This also correlates with centrosome amplification in cancer cells that have an overabundance of the long *CEP135* isoform relative to the short isoform (Ganapathi Sankaran et al., 2019). Alteration of this ratio affects centriole number. Furthermore, generation of the short mRNA isoform is cell cycle controlled (Dahl et al., 2015). Multiple genes important for cell cycle progression generally, and centriole assembly in particular, have been identified as alternatively spliced in a cell cycle dependent fashion. Much of this mRNA isoform control is tied to the CLK1 kinase which itself is cell cycle regulated by proteosomal degradation (Dominguez et al., 2016). The proper splicing of the γ-tubulin complex member, TUBGCP6, facilitated by WBP11, is required for centriole assembly (E. M. Park et al., 2020). Additionally, multiple mRNA splicing factors were identified in a genome-wide screen for genes required for centriole assembly and centriole number control (Balestra et al., 2013). Together, these examples point to the importance of transcript-level regulation in the control of centriole assembly. Yet, how specific splicing regulators affect centriole assembly remains poorly understood.

The multi-domained nuclear speckle and splicing factor protein, SON, is required for centriole assembly (Balestra et al., 2013; Ilik et al., 2020). SON is required for progression through the cell cycle and affects the efficient splicing of a number of genes important for microtubule nucleation and maintenance including γ-tubulin and Pericentrin (Ahn et al., 2011; Huen et al., 2010; Sharma et al., 2011). Loss of function mutations in SON are associated with intellectual disability, facial dysmorphology, scoliosis, heart defects and vision impairment, which are reminiscent of phenotypes associated with ciliopathies and could indicate defects in centriolar function (J.-H. Kim et al., 2016; Reiter & Leroux, 2017; Tokita et al., 2016).

Here we demonstrate that SON is required for centriole assembly, independent of its effects on the cell cycle. Amongst its many splicing targets, as revealed by mRNA-sequencing, is the gene encoding the centriolar satellite protein CEP131 whose reduction leads to severe loss of centriolar satellites. We find that centriolar satellites have an intimate relationship to assembling centrioles. Interestingly, *CEP131* has already been identified as a target of the splicing factor NRDE2 (Jiao et al., 2019), suggesting regulation of *CEP131* transcript formation has an important role in centriolar satellite formation and function. SON is required for the microtubule architecture around the centrosomes; its absence radically changes the number and distribution of microtubules around the centrosome and the makeup of the pericentrosomal region. This appears to be orchestrated by a transfer of microtubule nucleation activity from mother centrioles to their nascent daughters, facilitating the completion of procentriole assembly. This study highlights the importance of alternative splicing in modulating centriole assembly and microtubule organization, suggesting a multifactorial mechanism by which cells can modulate centrosome duplication, which is critical for cell division and development.

## Results

### The SON RNA splicing factor is required for centriole assembly

To explore the connection between centriole assembly and mRNA splicing, we selected five splicing factors from 14 that were identified in a global RNA interference screen for genes required for centriole assembly (Balestra et al., 2013). Some have roles in cell cycle progression ((SON and PRPF8); (Ahn et al., 2011; Wickramasinghe et al., 2015)), regulation of alternative splicing ((SON and PNN); (Chiu & Ouyang, 2006; Hickey et al., 2014)) and assembly of core splicing snRNP complexes ((SNRPD2 and SF3A3); (Siebring-van Olst et al., 2017; Tanackovic & Krämer, 2005; Urlaub et al., 2001)). We first asked whether knockdown of these splicing factors limited centriole assembly in a hTERT immortalized retinal epithelial cell line (RPE-1), which more closely mimics non-cancerous centriole assembly control, as they had in a cancer-derived cell line (HeLa) (Balestra et al., 2013). We employed an RPE-1 strain with GFP tagged *CETN2* to identify centrioles by centrin fluorescence, and PLK4 under control of the rtTA, doxycycline inducible promoter (Gossen et al., 1995; Hatch et al., 2010), and counted centrioles following 48 hours of splicing factor siRNA depletion (without PLK4 induction) (Figure 1A). Under these conditions, knockdown of SON, PNN and SNRPD2 demonstrated a significant decrease in the number of centrioles per cell, whereas knockdown of SF3A3 and PRPF8 had no observable effect on centriole frequency (Figure 1B).

**Figure 1.**
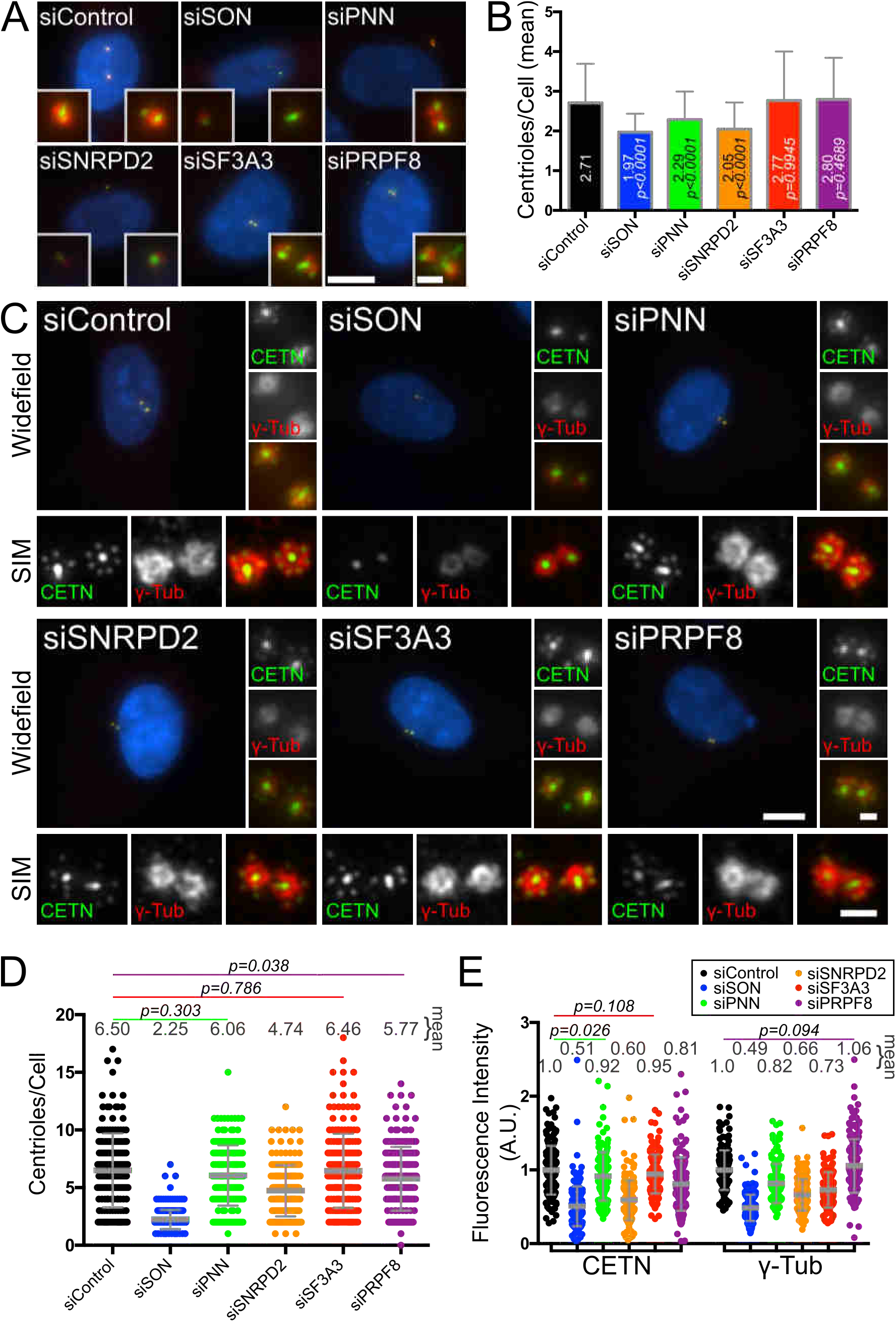
SON splicing factor depletion inhibits centriole assembly. A) SON depletion reduces centriole duplication in cycling cells. Cycling RPE-1/GFP-Cetn/Tet-PLK4 cells without PLK4 induction treated with the indicated siRNAs for 48 hours. Green, GFP-Cetn; Red, γ-Tubulin; Blue, DNA. B) Mean number of centrioles per cell. Mean and p values are indicated within the bars of the graph and p values are compared with siControl. N = siControl, 169 cells; siSON, 173 cells; siPNN, 173 cells; siSNRPD2, 169 cells; siSF3A3, 170 cells; siPRPF8, 180 cells. C) SON depletion reduces centriole duplication in S-phase arrested and PLK4 induced cells. siRNA-treated, S phase arrested cells induced for PLK4 expression. Top panels: widefield images; bottom panels: SIM images. D) The number of centrioles per cell after S phase arrest and PLK4 induction. p values are < 0.0001 unless otherwise indicated. E) Fluorescence quantification of GFP-Cetn and γ-tubulin after S phase arrest and PLK4 induction. p values are < 0.0001 unless otherwise indicated. Scale bars for whole cells = 10 μm, for centrosomes = 1 μm. N for D) and E) = siControl, 177 cells; siSON, 204 cells; siPNN, 194 cells; siSNRPD2, 189 cells; siSF3A3, 195 cells; siPRPF8, 203 cells. Data compiled from two biological replicates. Bar and error bars: mean and standard deviation.

Splicing factors commonly have pleiotropic effects and often effect the cell cycle (Abramczuk et al., 2017; Ahn et al., 2011; Kurokawa et al., 2014; Suvorova et al., 2013). Centriole assembly is intimately linked to the cell cycle, and thus alterations to the cell cycle could affect centriole numbers in a manner indirectly related to centriole assembly control. To address this, we isolated centriole assembly from other processes by observing cells under conditions of S phase arrest with overexpression of PLK4 to promote centriole assembly (Figure 1C) (Kleylein-Sohn et al., 2007). In 72% of cells, PLK4 induction results in procentriole assembly around the mother centriole, often in a rosette formation (Kleylein-Sohn et al., 2007). Centrin foci counts of cells under these conditions revealed that knockdown of SON and SNRPD2 resulted in significant reductions to centriole number when cell cycle effects were minimized. SON knockdown had by far the strongest effect, as only 2% of cells have procentrioles, based on Centrin labeling (Figure 1D). Fluorescence quantification of Centrin showed reduced levels with SON knockdown, commensurate with the reduction in centriole number (Figure 1E). Centrin was also reduced with SNRPD2 knockdown and to a lesser extent with PRPF8 knockdown, suggesting that these proteins affect the amount of Centrin protein present in daughter centrioles. Quantification of γ-tubulin fluorescence intensity did not completely correlate with Centrin fluorescence, suggesting that different splicing factors affect centrosomal proteins in distinct ways (Figure 1E, see siSF3A3 and siPRPF8). Consistent with the reduction in γ-tubulin observed with SON knockdown, SON has previously been shown to be a splicing factor important for intron removal in *TUBG1* (γ-tubulin encoding gene) and affects γ-tubulin protein levels (Ahn et al., 2011). We confirmed the dramatic impact on daughter centriole assembly in SON knockdown with a second siRNA which had similar efficacy and effects on centriole assembly (Figure Supplement 1A, B). Furthermore, S phase arrest in *SON* knockdown is similar to controls (Figure Supplement 1C). Because depletion of SON reduces centriole assembly independent of the cell cycle, we focused on this splicing factor to identify what steps of centriole assembly are disrupted.

### SON depletion is permissive for early steps of centriole assembly

To determine whether the initiating events of centriole assembly are affected when SON is depleted, we examined the localization of core centriole assembly factors. CEP152 and CEP192 are centriole scaffold proteins that interact with and are required for PLK4 recruitment to the centriole (T.-S. Kim et al., 2013; Sonnen et al., 2013). Total CEP152 and CEP192 are reduced at the centrosome in siSON (59% of siControl for both), but the mother centriole levels are only reduced to 75% of controls, suggesting that some of the reduction in protein levels is due to a lack of recruitment to procentrioles (Figure 2A, B, Figure Supplement 2B). Despite reductions of these scaffold proteins, levels of PLK4 at the centrosome are unaffected, suggesting that the defect in centriole assembly is not due to a failure to recruit PLK4 (Figure 2A, B).

**Figure 2.**
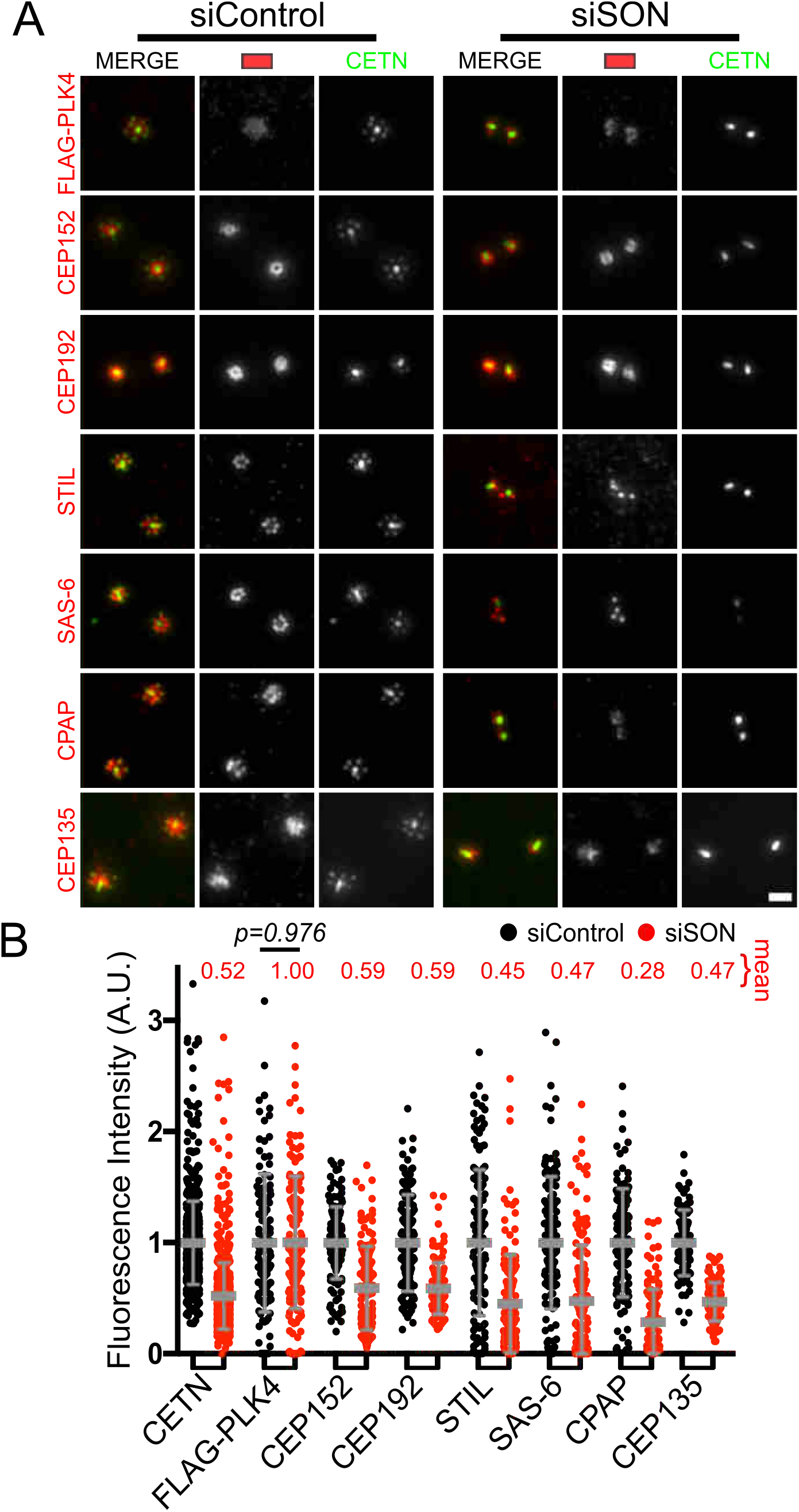
SON depletion reduces but does not eliminate early centriole assembly. A) Structured Illumination Microscopy (SIM) images of siRNA treated, S phase arrested cells induced for PLK4 expression and stained for centriole assembly factors. Scale bar = 1 μm. B) Fluorescence intensity quantifications of widefield microscopy images (Supplementary Figureure 2A). N = CETN: siControl, 851 centrosomes; siSON, 925 centrosomes; FLAG-PLK4: siControl, 111 centrosomes; siSON, 131 centrosomes; CEP152: siControl, 136 centrosomes; siSON, 143 centrosomes; CEP192: siControl, 124 centrosomes; siSON, 123 centrosomes; STIL: siControl, 136 centrosomes; siSON 138 centrosomes; SAS-6: siControl, 132 centrosomes; siSON, 148 centrosomes; CPAP: siControl, 120 centrosomes; siSON, 122 centrosomes; Cep135: siControl, 91 centrosomes; siSON, 119 centrosomes. Data compiled from two biological replicates. Bar and error bars: mean and standard deviation.

Assembly of new centrioles begins with STIL phosphorylation by PLK4, allowing for SAS-6 recruitment to nascent daughter centrioles where it forms the inner cartwheel which is an early assembly intermediate in centriole assembly (Tyler Chistopher Moyer & Holland, 2019; Ohta et al., 2014). CPAP and CEP135 promote the connection of the cartwheel to the microtubule triplet walls of the growing centriole (Y.-C. Lin et al., 2013). We next asked whether the levels of these early assembly factors are altered upon SON depletion, and if so, whether these alterations could result in the observed centriole assembly defect. Although the mean levels of these early assembly factors are reduced in siSON (STIL = 45%, SAS-6 = 47%, CPAP = 28% and CEP135 = 47%), in no case did the abundance of any of these proteins correlate with more centriole assembly in individual cells in the siSON condition (Figure Supplement 2A, C). As expected in siControl treated cells, both STIL and SAS-6 protein levels correlate with the number of centrioles as overexpression of either is sufficient to promote centriole assembly (Figure Supplement 2C) (Leidel et al., 2005; Vulprecht et al., 2012). These data indicate that early centriole assembly factors are reduced but not eliminated during SON depletion. Furthermore, because a sufficient amount of these assembly factors in individual siSON cells does not correlate with more centrioles, these data suggest that their overall decrease in abundance is not responsible for the centriole assembly phenotype.

To ascertain whether centriole assembly initiates when SON is depleted, we examined SAS-6 levels. Because SAS-6 is lost from mature centrioles during exit from mitosis, SAS-6 signal exclusively marks assembling procentrioles (Puklowski et al., 2011). When PLK4 is induced 86.4% of siControl-treated cells have three or more centrioles based on centrin foci indicating new centriole assembly (Figure Supplement 3). This allowed us to set a SAS-6 fluorescence intensity cut-off for new centriole assembly. 56% of siSON cells had a SAS-6 signal above this threshold. This indicates that more than half the SON depleted cells have initiated centriole assembly (Figure Supplement 3). We then utilized structured illumination microscopy (SIM) to examine siSON cells to determine whether early assembly factors were indeed present in a rosette conFigureuration consistent with early procentriole assembly. Rosettes of STIL, SAS-6, CPAP and CEP135 were detected around mother centrioles, even though no Centrin labeled procentriolar foci were observed (Figure 3A). This indicates that centriole assembly initiates in SON knockdown cells but is unable to proceed to completion.

**Figure 3.**
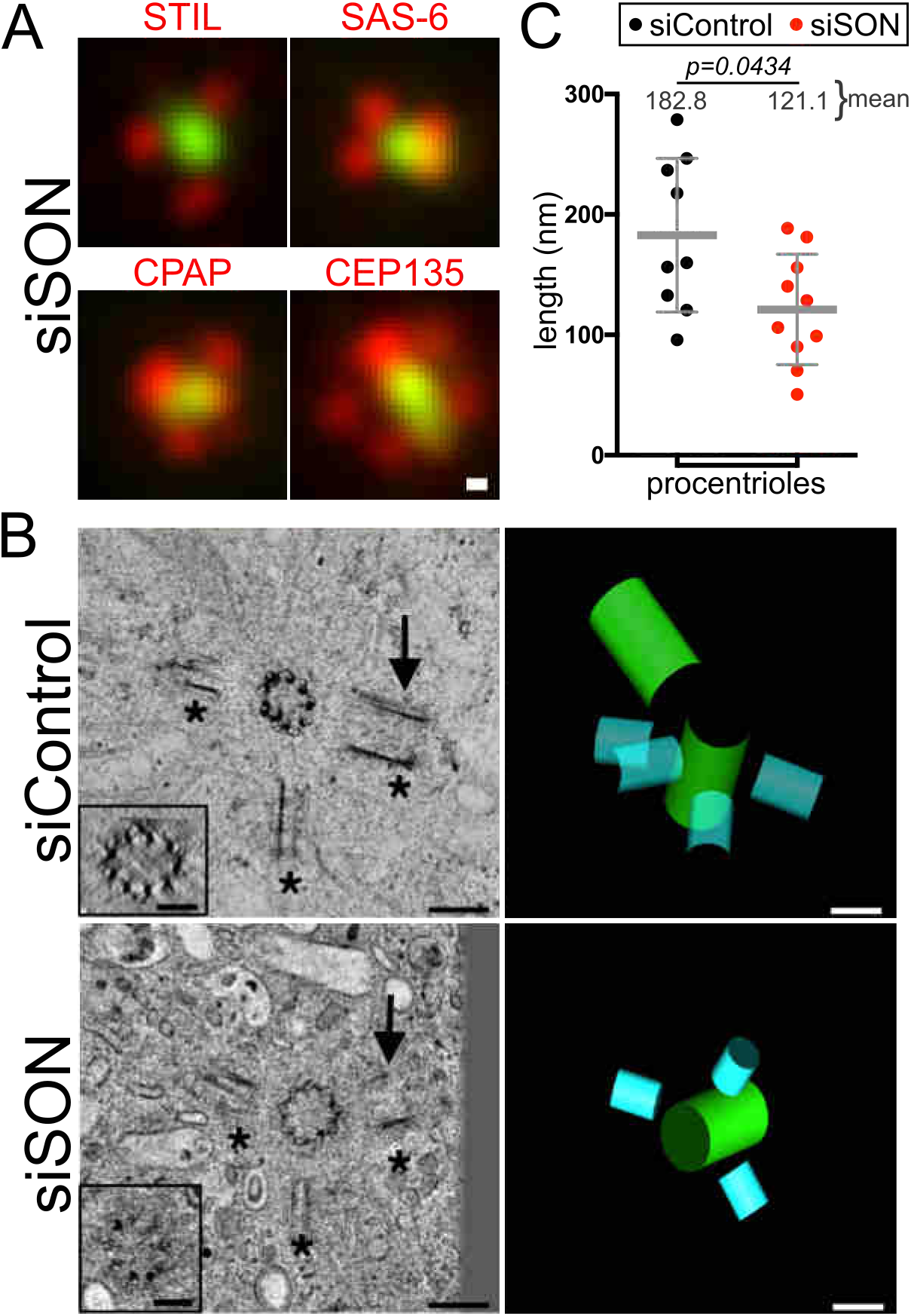
SON depletion causes short and immature procentrioles with missing C-tubules. A) SIM images of siSON treated cells stained as indicated showing that proteins required for early steps of procentriole assembly are present. Scale bar = 100 nm. B) Tomographic slices and the corresponding tomographic models from siControl and siSON treated cells. Procentrioles are present in both conditions and represented in blue in the models (asterisks indicate 3 of the 4 procentrioles in siControl that are visible in this view). The arrows indicate the position of procentrioles that were rotated to reveal microtubule triplets in the control and microtubule doublets when SON is depleted (insets). The A-tubules are densely stained. The 3D model of the siControl rosette shows two mature centrioles (green) and Movie S1 shows the complete volume, built from four 250 nm thick sections. The siSON centrosomes are separated and the adjacent centrosome is presented in Figure Supplement 7A (D’’). Movie S2 shows the complete volume, which contains both centrosomes, and is built from four 250 nm thick sections. Scale bars = 200 nm and 100 nm for insets. C) The length of procentrioles in tomographic volumes. N = siControl, 9 procentrioles; siSON, 10 procentrioles. Bar and error bars: mean and standard deviation.

To determine the structural events associated with these new procentriole assemblies, we used electron tomography. Consistent with the immunofluorescence data, when SON is depleted, three out of four cells examined exhibited incomplete procentriole rosettes with procentrioles that were on average shorter than those observed in control cells (Figure 3B, C). Moreover, the microtubules in these procentrioles lack the C-tubule which is present in control conditions (Figure 3B insets, Movie S1, Movie S2). We therefore conclude that SON depletion prevents daughter centriole assembly at a stage after early procentriole assembly and initial microtubule nucleation events, but prior to complete triplet microtubule formation, Centrin deposition and centriole elongation.

### SON impacts splicing of genes encoding centriolar satellite and microtubule proteins

To determine SON splicing targets responsible for the centriole assembly defect, we utilized global mRNA sequencing of RNA isolated from cells under conditions of S phase arrest and PLK4 induction. From these data we identified 4,413 genes downregulated in SON depleted cells (Table S1). To establish whether SON has a greater effect on specific cellular structures, we identified Cellular Component (CC) Gene Ontology (GO) terms most effected by SON depletion. The most highly enriched GO terms include the microtubule cytoskeleton and centrosome (Figure 4A, Table S1), which is consistent with the microtubule disruption previously observed after SON knockdown (Ahn et al., 2011; Sharma et al., 2011). Along with changes to expression levels, we also examined changes in alternative splicing using the software packages MAJIQ (Table S2) (Vaquero-Garcia et al., 2016) and rMATS (Table S3) (Shen et al., 2014). From this analysis we found 2,996 genes by MAJIQ and 3,531 genes by rMATS that showed significant changes in alternative splicing (Figure 4B). These two software packages identified genes with considerable overlap (74.3% of MAJIQ identified genes in the rMATS set and 63.1% rMATS identified genes in the MAJIQ set, Figure Supplement 4A). These genes also showed strong enrichment for microtubule-associated GO terms including microtubule cytoskeleton, microtubule organizing center, centrosome and centriole (Figure 4A). To investigate additional targets of SON that could contribute to the centriole assembly defect, we also tested for overlap with genes included in the Centrosome Database (CD, portion of the Centrosome and Cilia Database (Gupta et al., 2015)) and genes encoding proteins identified in an extensive proximity mapping of centriolar satellites (Gheiratmand et al., 2019). Interestingly, differentially spliced genes strongly overlapped with both of these additional lists, indicating that these proteins are affected at a high rate by SON knockdown (Figure 4A, B).

**Figure 4.**
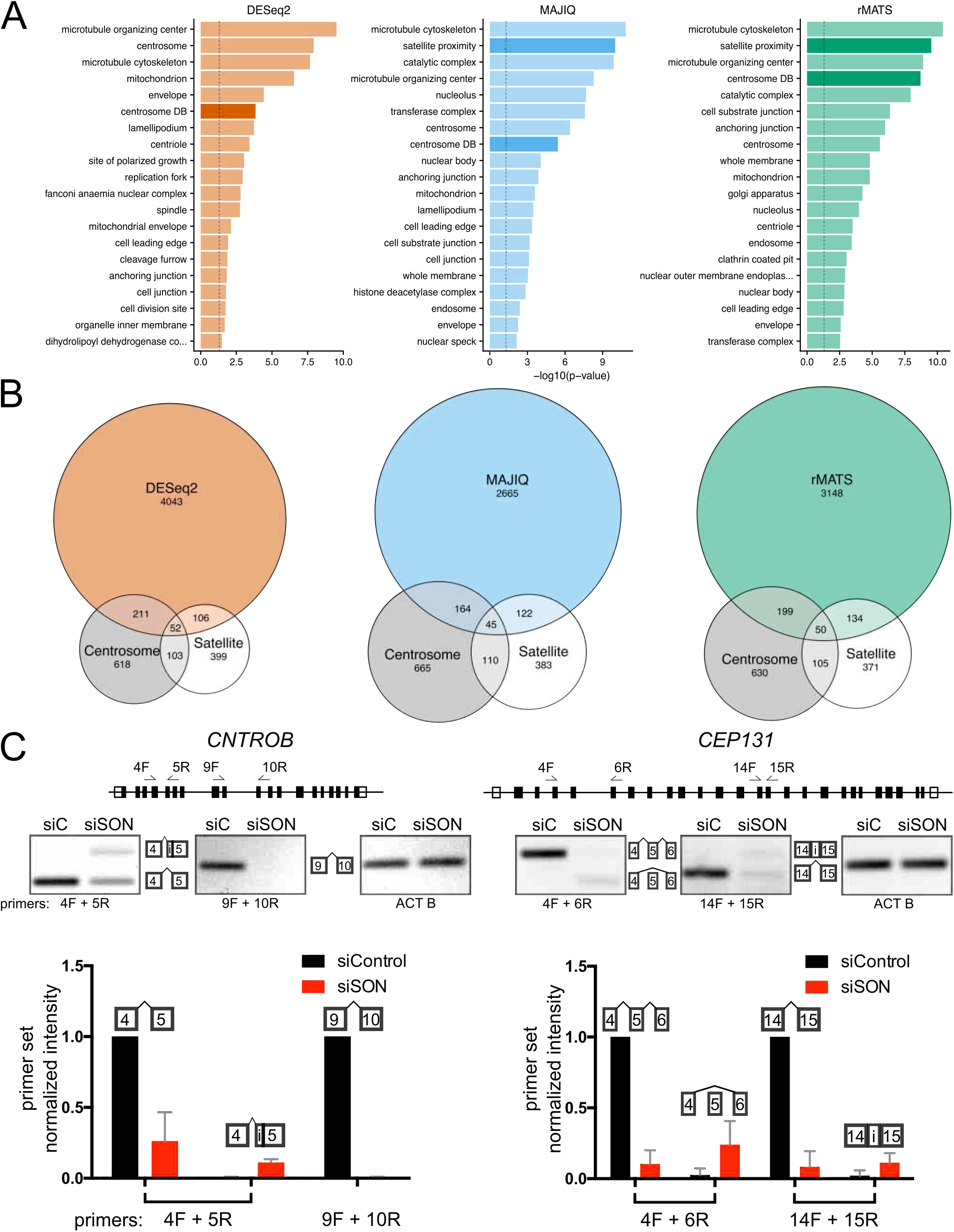
SON depletion disrupts splicing of genes responsible for centriole assembly, centriolar satellites and the microtubule cytoskeleton. A) The centrosome fraction of the Centrosome and Cilia Database (CCDB, centrosome DB) (Gupta et al., 2015) and genes determined by proximity mapping of centriolar satellites (satellite proximity) (Gheiratmand et al., 2019) were included with Cellular Component Gene Ontology (GO) terms in opaque colors. The top 20 terms for reduced gene expression (DESeq2) and alternative splicing (as determined by either the MAJIQ or rMATS software packages) are ordered by p values. The dashed lines indicate a p value of 0.05. B) Venn diagrams displaying the overlap of the centrosome database (Centrosome), the centriolar satellite proximity list (Satellite) and reduced expression or spliced genes. Areas of each circle are proportionate with the gene set. C) SON depletion results in improper splicing of *CNTROB* and *CEP131*. Top panels show the location of introns and exons and the primers used for reverse transcriptase PCR (RT-PCR), middle panel. Bottom panel: quantification of the RT-PCR bands normalized first to the amount of RNA as determined by ACT B RT-PCR and second to the amount of the properly spliced fragment in the control condition. RNA-seq and RT-PCR were performed and quantified on three biological replicates. Error bars: standard deviation.

Genes from the centrosome database and satellite proximity lists whose splicing is affected by SON were then sorted by their differential expression (Table S4). Within the top 50 genes in this list were several with known functions at centrosomes, including *TUBG1* (γ-tubulin), *PCNT* (Pericentrin), *CEP131*, *HAUS4* (augmin complex member) and *CNTROB* (Centrobin) (Delaval & Doxsey, 2010; Staples et al., 2012; Thawani et al., 2019; Zou et al., 2005). Intron excision events in *TUBG1* and *PCNT* had previously been reported as defective with SON depletion (Ahn et al., 2011), and splicing changes to *CEP131* were detected in a micro array analysis (Sharma et al., 2011). To verify changes to *CNTROB* and *CEP131* splicing upon SON reduction, reverse transcription followed by PCR was performed. In addition to reducing the amount of *CNTROB* and *CEP131* mRNA, changes in alternative splicing events were observed including mutually exclusive exons, alternative splice site usage, and exon skipping (Figure 4C, Figure Supplement 4C, Tables S2 and S3). Our RNA sequencing data confirms *TUBG1* and *PCNT* require SON for correct splicing (Figure Supplement 4C) and identifies *CNTROB* and *CEP131* as additional targets that could impact centriole assembly. Therefore, the SON splicing factor is required for the homeostatic control of mRNAs encoding centrosomal, centriolar satellite and microtubule cytoskeletal proteins.

### SON depletion reduces centriolar satellites and trafficking structures

We examined the distributions of centrosomal and centriolar satellite proteins whose splicing is affected by SON using a semi-automated radial intensity analysis algorithm in control and SON knockdown cells (Figure 5A, B) (Sankaran et al., 2020). γ-tubulin, together with γ-tubulin complex members, acts to nucleate microtubules and is concentrated at the centrosome (Stearns et al., 1991; Tovey & Conduit, 2018). SON depletion did not alter γ-tubulin fluorescence intensity at the centrosome itself. However, a significant reduction in γ-tubulin was observed in the region 1.5 to 2.5 µm from the centrosome. SON knockdown had a similar effect on the distribution of Pericentrin, a large scaffolding protein that interacts with γ-tubulin (Dictenberg et al., 1998) and localizes both to the centrosome and to mobile cytoplasmic granules that transit to and from the centrosome (Galati et al., 2018; Young et al., 2000). The Pericentrin in cytoplasmic granules is nearly eliminated, suggesting that Pericentrin-dependent trafficking to and from centrosomes is severely affected when SON is depleted (Figure 5A, B). Centrobin is associated with daughter centrioles and important for their elongation (Gudi et al., 2011; Zou et al., 2005). Centrobin also limits the recruitment of additional PCM proteins (Jeffery et al., 2013), stabilizes CPAP (Gudi et al., 2014, 2015) and promotes C-tubule formation and maintenance (Reina et al., 2018). This makes Centrobin an intriguing candidate for the SON knockdown centriole assembly phenotype as cells with reduced SON initiate but do not complete centriole assembly (Figure 3A, B), have a larger decrease in centrosomal CPAP than other early centriole assembly factors (Figure 2B), and produce procentrioles without the C-tubule (Figure 3B). We observed that Centrobin fluorescence intensity is reduced at the centrosome in the SON knockdown (Figure 5A, B). CEP131 is an integral component of centriolar satellites and important for cilia formation and genomic stability (Denu et al., 2019; Graser et al., 2007; Staples et al., 2012). CEP131 has a significant reduction both near the centrosome and in the region surrounding the centrosome. Measuring total fluorescence within a 5 μm radius around the centrosome shows reductions to the mean of 51%, 41%, 33% and 16% of control levels for γ-Tubulin, Centrobin, Pericentrin and CEP131, respectively (Figure Supplement 5B).

**Figure 5.**
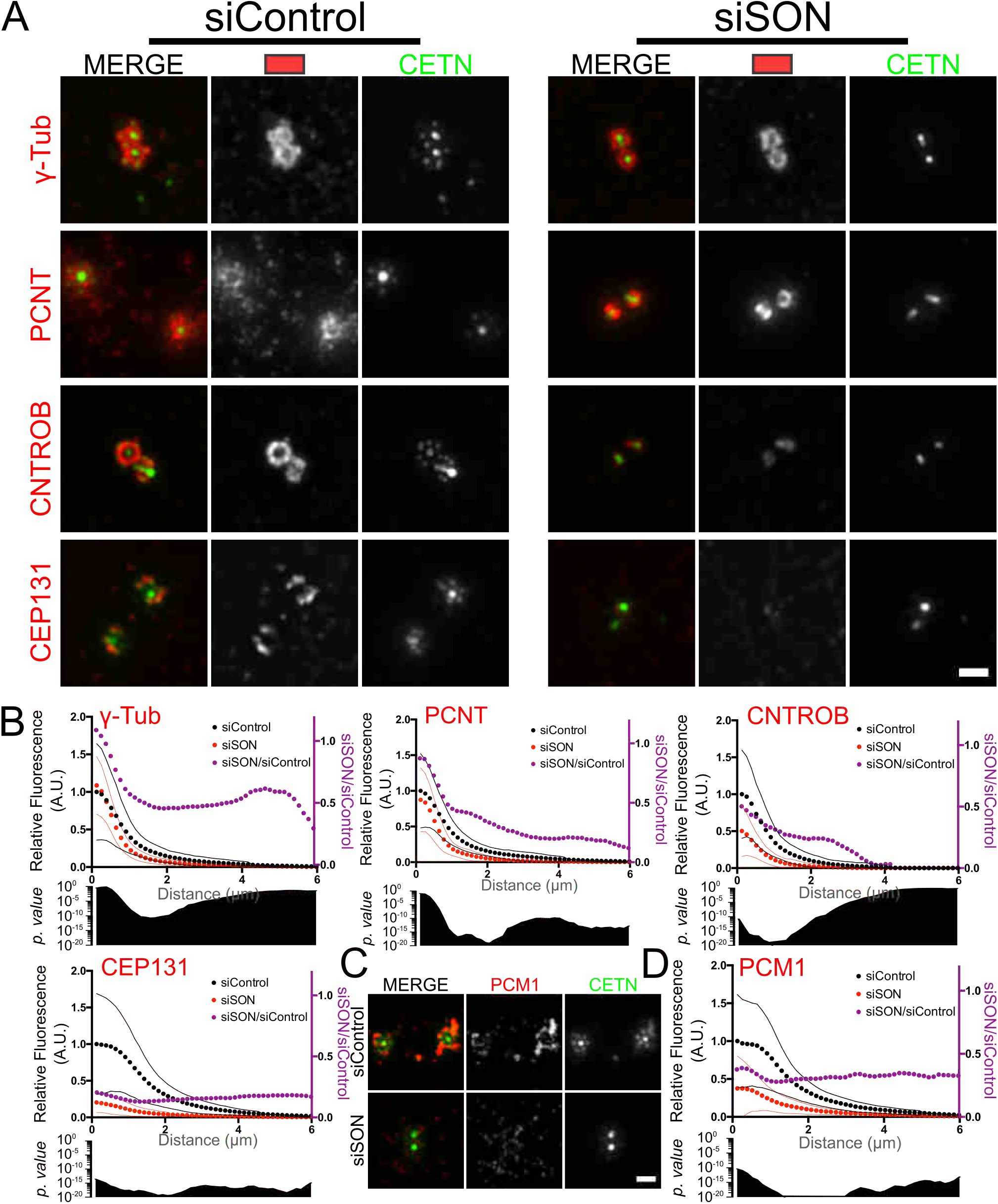
SON splicing targets disrupt Centrobin, centriolar satellites and trafficking granules. A) SON targets are diminished at the centrosome. SIM images of siRNA treated, S phase arrested cells induced for PLK4 expression and stained for the identified SON targets. Scale bar = 1 μm. B) γ-tubulin and Pericentrin are reduced most in the region 1.5 to 2.5 µm from the centrosome center. Normalized mean radial quantification of fluorescence intensities from widefield images (Figure Supplement 5A) (Y axis on left, black = siControl, red = siSON). The ratio of the intensity of siSON/siControl is also plotted (Y axis on right, purple). p values from the Student’s T-test are plotted beneath each graph on a semi-log scale. The mean value is plotted, and standard deviations are plotted as thin lines in the corresponding color. N = γ- Tubulin: siControl, 76 centrosomes; siSON, 73 centrosomes; Pericentrin (PCNT): siControl, 72 centrosomes; siSON 76 centrosomes; Centrobin (CNTROB): siControl, 101 centrosomes; siSON, 106 centrosomes; CEP131: siControl, 70 centrosomes; siSON, 96 centrosomes. Data are compiled from two biological replicates. C) SIM images of siRNA treated, S phase arrested cells induced for PLK4 expression and stained for PCM1. Scale bar = 1 μm. D) Radial quantification of PCM1 fluorescence intensity as in B). N = siControl, 78 centrosomes; siSON, 70 centrosomes. Data is compiled from two biological replicates.

To understand which changes to protein levels might explain the centriole assembly defect in SON knockdown, we next measured the fluorescence levels of centrosome localized γ-tubulin, Centrobin, Pericentrin and CEP131 relative to the frequency of centriole assembly. Correlating fluorescence at and around the centrosome with centriole counts suggested that γ-tubulin levels at and near the centrosome do not explain the SON depletion centriole assembly defect because individual cells with γ-tubulin at levels similar to control cells do not have more centrioles. Importantly, Centrobin, Pericentrin and CEP131 were reduced to levels in SON knockdown that suggest they contribute to the centriole assembly phenotype (Figure Supplement 5C).

Because reduction of CEP131 and Pericentrin indicated profound changes to the trafficking structures and centriolar satellites around the centrosome, we investigated centriolar satellites further. We examined localization of the canonical centriolar satellite scaffold protein, PCM1, in the SON knockdown even though siSON’s effect on *PCM1* mRNA expression was modest (85% of control). PCM1 and CEP131 are both major centriolar satellite proteins with considerable overlap in their localization at centriolar satellites (Dammermann & Merdes, 2002; Gheiratmand et al., 2019; Kubo et al., 1999; Prosser & Pelletier, 2020; Staples et al., 2012). Similar to CEP131, PCM1 fluorescence intensity was reduced throughout the radial analysis, although not to the same degree (34% of control versus 16% for CEP131) (Figure 5C, D and Figure Supplement 5A, B). Taken together, this indicates that SON is required for the proper levels and distribution of centriolar satellites.

To assess whether the reduction in centriolar satellite fluorescence intensities at or around the centrosomes was due to mis-localization or total cellular loss of protein, total fluorescence levels inside cells were quantified in both siSON and siControl cells (Figure Supplement 5D). Some proteins whose genes depend upon SON for correct splicing had significant reductions in overall protein levels (83% for γ-tubulin, 53% for Pericentrin and 44% for CEP131). PCM1 was also reduced in total protein levels (78%) but alternative splicing was not significantly affected by SON depletion, suggesting that when satellites are disrupted, PCM1 protein stability is affected. We also detected an elevation in total cellular SAS-6 protein levels, even though there was not a large transcriptional change as determined by RNA-sequencing (1.06 fold change relative to siControl). In summary, SON activity is required for the normal accumulation and distribution of centriolar satellites and Pericentrin trafficking structures. It may do so through its robust effect on CEP131 protein levels.

SIM imaging revealed localization of PCM1 and CEP131 directly adjacent to nascent procentrioles in siControl cells (Figure 5A, C). This localization, as well as the formation of Centrin positive procentrioles, is ablated when SON is depleted. This suggests that centriolar satellites (which are also important for Centrin accumulation at centrosomes (Dammermann & Merdes, 2002)) and trafficking particles participate in late steps of centriole assembly and are affected by SON.

### Depletion of centrosome and centriolar satellite targets of SON phenocopies SON depletion

To assess the contributions of the SON targets Centrobin, CEP131 and Pericentrin to the centriole assembly defect observed in SON depletion, we evaluated centriole assembly upon knockdown of these individual components. Knockdowns of *CNTROB* and *CEP131* mRNAs effectively reduced the corresponding proteins at centrosomes in the PLK4 overexpression cells (Figure 6A, B). However, centriole assembly was only modestly affected upon knockdown of Centrobin or CEP131 individually or in combination (Figure 6C), indicating that these proteins, on their own, do not account for the reduction in centriole assembly observed with SON knockdown. Because Centrobin has been reported to stabilize CPAP (Gudi et al., 2014, 2015), we examined whether Centrobin knockdown would recapitulate the reduction in CPAP observed when SON is depleted (Figure 2B). Surprisingly, Centrobin depletion resulted in a modest increase to CPAP fluorescence intensities at centrosomes in the PLK4 induction experiments (Figure Supplement 6A, B). Therefore, splicing changes to Centrobin as a result of SON depletion are unlikely to contribute to the reduced CPAP levels and centriole assembly observed in the SON knockdown.

**Figure 6.**
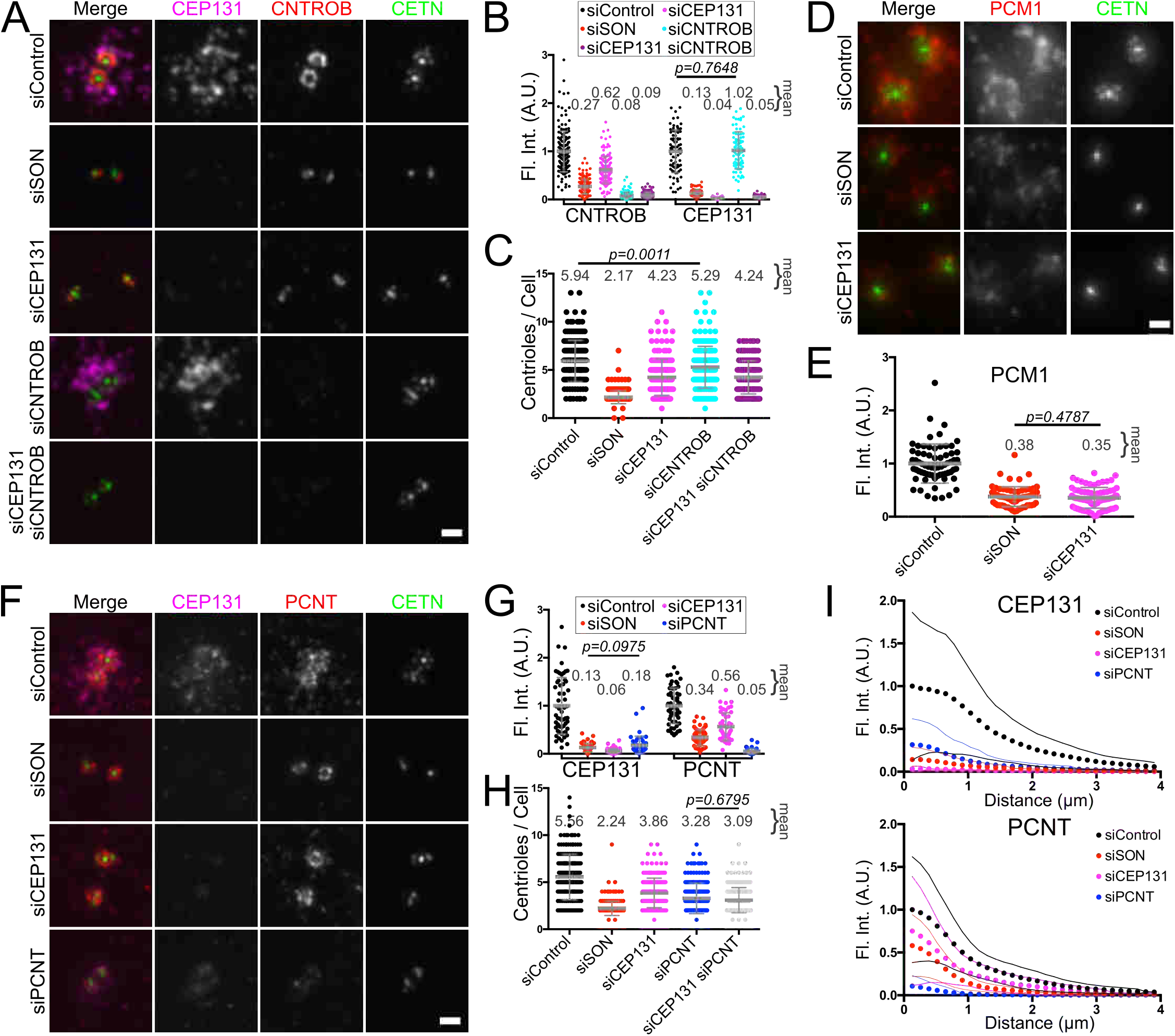
Depletion of SON splicing targets partially phenocopy SON depletion. A) *CEP131* knockdown partially inhibits centriole assembly. SIM images of CEP131 and Centrobin (CNTROB) stained cells depleted of SON, CEP131, Centrobin, or both CEP131 and Centrobin. B) Fluorescence quantification of widefield images of CEP131 (5 μm radius - cell) and Centrobin (1 μm radius - centrosome). N = siControl: 165 centrosomes, 85 cells; siSON: 158 centrosomes, 78 cells; siCEP131: 151 centrosomes, 71 cells; siCNTROB: 161 centrosomes, 81 cells; siCEP131 and siCNTROB: 173 centrosomes, 86 cells. C) Centriole counts after S phase arrest and PLK4 induction in indicated siRNA conditions. N = 180 cells for each condition. Quantifications are from two biological replicates. D) Depletion of CEP131 results in a reduction in PCM1 around the centrosome. Widefield images of cells stained for PCM1 after SON or CEP131 depletion and S phase arrest and PLK4 induction. E) PCM1 fluorescence quantification of a 5 μm radius circle around centrosomes in images from D). N = siControl: 67 cells; siSON: 68 cells; siCEP131: 75 cells. Quantifications are from two biological replicates. F) Pericentrin depletion reduces centriole assembly and CEP131 localization. SIM images of S phase arrested and PLK4 induced cells stained for CEP131 and Pericentrin (PCNT). SON, CEP131 and Pericentrin (PCNT) were depleted as indicated. G) Fluorescence quantification of widefield images of CEP131 and Pericentrin (PCNT) within a 5 μm radius around the centrosomes. N = siControl: 57 cells; siSON: 56 cells; siCEP131: 49 cells; siPCNT: 45 cells from two biological replicates. H) Centriole counts after S phase arrest and PLK4 induction in indicated siRNA conditions. N = 186 cells for each condition. Quantifications are from two biological replicates. I) Mean radial fluorescence intensity measurements from the centrosome of CEP131 and Pericentrin (PCNT). The standard deviation is plotted as thin lines in the corresponding colors. N = siControl: 63 cells; siSON: 61 cells; siCEP131: 46 cells; siPCNT: 44 cells. Quantifications are from two biological replicates. Bar and error bars: mean and standard deviation. p values are < 0.0001 unless otherwise displayed. Scale bars = 1 μm.

Because Centrobin had a modest effect on centriole assembly, we focused on the effects of CEP131 and Pericentrin. Localization of PCM1 in both SON depletion and CEP131 depletion was reduced to similar levels (Figure 6D, E), indicating that the PCM1 reduction observed with SON depletion can be attributed to the reduction in CEP131. CEP131 has also been implicated in the trafficking of CEP152 to the centrosome (Denu et al., 2019; Kodani et al., 2015). To assess whether the centrosomal CEP152 reduction observed with SON depletion is due to loss of CEP131, we quantified centrosomal CEP152 fluorescence in both SON and CEP131 depletion conditions. Reductions in CEP152 were observed in both conditions (Figure Supplement 6C, D). Treatment with siCEP131 was more effective at reducing CEP131 levels than treatment with siSON, but CEP152 levels in siSON were reduced by slightly more than by siCEP131 alone. This suggests that promotion of correct *CEP131* splicing by SON is partially responsible for the observed reduction in CEP152.

We next examined how Pericentrin depletion in siSON contributes to the centriole assembly defect. Depletion of Pericentrin was efficient, and reduced centriole assembly to a greater degree than depletion of CEP131, although not to the level of SON depletion (Figure 6F, G, H). Depletion of both CEP131 and Pericentrin concurrently had a similar effect on centriole assembly as depletion of Pericentrin alone (Figure 6H). SON depletion, interestingly, reduces Pericentrin directly surrounding the mother centrioles at the centrosomes to 60-80% of controls (Figures 5A, B, 6F, I) whereas Pericentrin depletion reduces this population to 15% (Figure 6F, I), making the contribution of SON dependent splicing of Pericentrin to the centriole assembly defect difficult to assess (Figure 6F, I). However, it raises the possibility that SON is specifically necessary for the trafficking population of Pericentrin (Figure 5A, B). Additionally, we examined the interplay between CEP131 and Pericentrin. Consistent with previous reports (Staples et al., 2012), CEP131 depletion results in a modest decrease in Pericentrin fluorescence intensity around the centrosome (56%), while Pericentrin depletion causes a dramatic decrease in CEP131 fluorescence intensity around the centrosome (18%) (Figure 6G, I). Therefore, reductions in the SON splicing targets CEP131 and Pericentrin account for the loss of centriolar satellites observed in cells depleted of SON, but do not fully account for the SON depletion centriolar assembly defect.

Because SON depletion is permissive for early steps of centriole assembly but prevents advancement of the assembly process, we examined CEP131 and Pericentrin depletion for procentriole intermediates as found in the SON knockdown. We utilized SAS-6 fluorescence as a proxy for these procentriole intermediates. In SON depleted cells, we once again observed SAS-6 signal in a rosette formation without Centrin foci. Early centriole assembly intermediates were never observed when Pericentrin was depleted, which is consistent with reports that SAS-6 requires Pericentrin for its localization (Ito et al., 2019), and only rarely observed when CEP131 was depleted (Figure Supplement 6E, F, G). We therefore conclude that when SAS-6 is present at procentrioles in Pericentrin depleted cells, centriole assembly is able to proceed. When SAS-6 is present at procentrioles in CEP131 depleted cells, centriole assembly usually proceeds, but occasionally is slowed or arrested, as is the case for most cells depleted of SON (Figure Supplement 6G). Given that CEP131 depletion most closely mimics SON depletion, we asked whether *CEP131* could rescue the centriole assembly defect in SON depleted cells. *CEP131* was unable to rescue the SON depletion phenotype, and overexpression of exogenous *CEP131* inhibited centriole assembly (Figure Supplement 6H, I).

In total, CEP131 and Pericentrin depletion can recapitulate some of the centriole assembly defects observed when SON is depleted, but depletion of these proteins individually or in combination, nor can depletion of Centrobin, explain the severe centriole assembly defect observed upon SON depletion, suggesting the mechanism of SON-based regulation of centriole assembly is multifactorial.

### SON depletion alters the microtubule landscape near the centrosome

Because centriolar satellites utilize microtubule-dependent trafficking (Conkar et al., 2019; Kubo et al., 1999), and because we observed close association between centriolar satellites and the assembling procentrioles, we examined the microtubule network near centrosomes. We observed a 2-fold increase in microtubule density in SON depleted cells, consistent with previous reports (Figure 7A, B, Figure Supplement 4A, B, C) (Ahn et al., 2011). Using SIM, we resolved the relationship between assembling centrioles, microtubules and centriolar satellites (Figure 7C). Microtubules originated from nascent procentrioles. Furthermore, the centriolar satellite protein, PCM1, is positioned at the ends of these microtubules, suggesting that assembling centrioles have microtubules and centriolar satellites in close proximity to provide components required for their elongation and maturation. When SON is depleted, the microtubule arrangement is distinctly different, with microtubules crowded around mother centrioles. To quantify this rearrangement, we examined the microtubule intensity around mother centrioles to determine the average distance at which microtubules begin. Because the average intensity of microtubules dissipates as a function of distance from the centrosome (Figure 7B), we examined the shift from increasing microtubule intensity to decreasing intensity as measurements are quantified radially from the mother centriole as a way to establish the boundary of microtubule minus-ends close to the centrosome. This occurred on average 480 nm from the mother centrioles in siControl cells and 340 nm in siSON cells (Figure 7C, D) indicating that not only are there more microtubules around centrosomes when SON is depleted but that the network encroaches closer to the mother centrioles.

**Figure 7.**
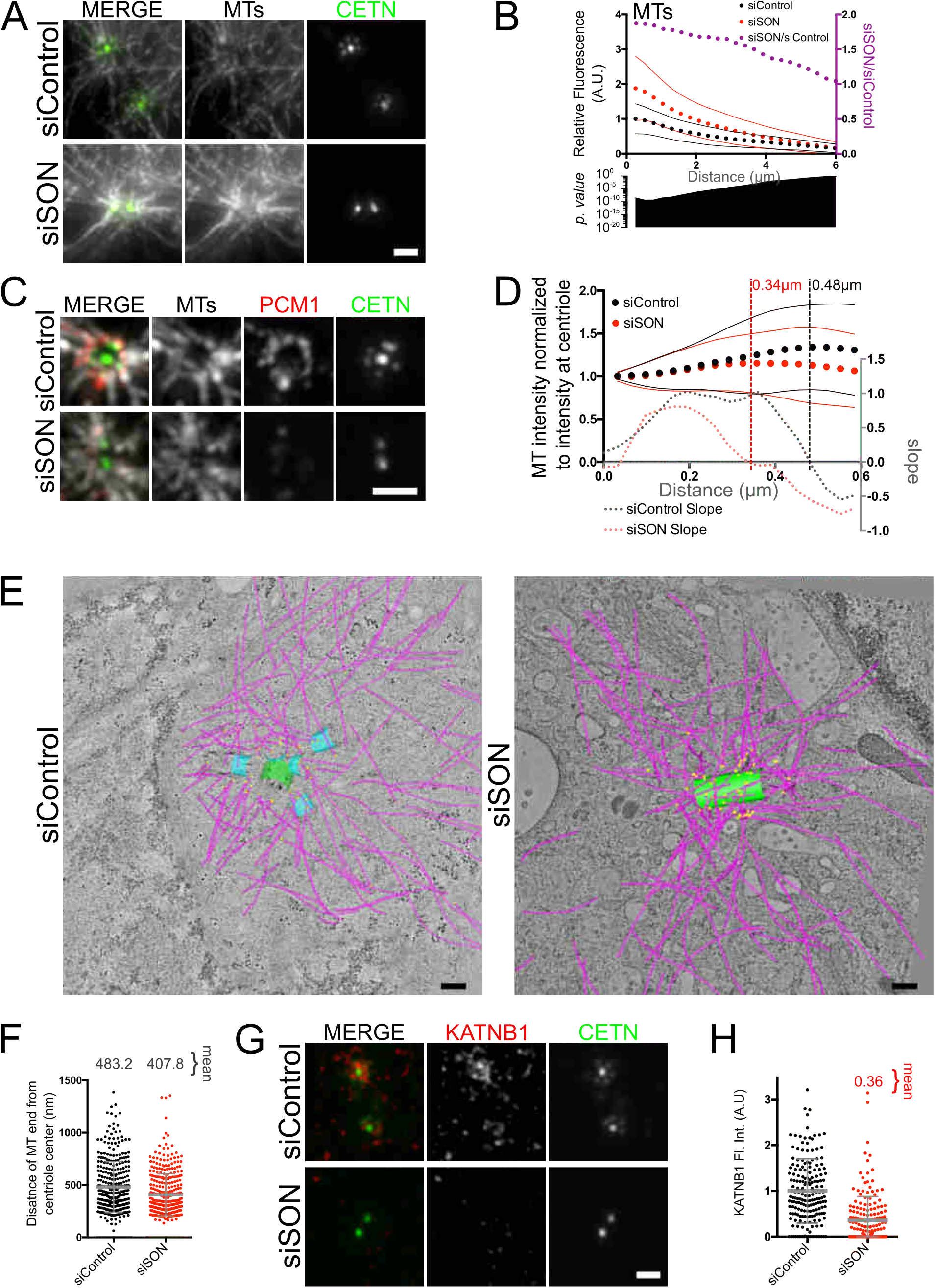
SON depletion increases the density of centrosomal microtubules from the mother centriole and reduces Katanin at the centrosome. A) The density of microtubules around the centrosome is increased when SON is depleted. SIM images of siRNA treated, S phase arrested cells induced for PLK4 expression and stained for microtubules (gray). B) Radial quantification of microtubule fluorescence intensities from the centrosome. N = siControl: 46 cells; siSON: 63 cells from two biological replicates. C) Procentriole, centriolar satellite and microtubule colocalization is disrupted by SON depletion. SIM images of siRNA treated, S phase arrested cells induced for PLK4 expression and stained for PCM1 (Red) and microtubules (gray). D) Microtubules originate closer to the mother centriole in SON depleted cells. Microtubule fluorescence intensities from line scans of SIM images of ten mother centrioles (five cells) for both siControl and siSON (Y axis on left, black = siControl, red = siSON). Intensities were normalized to the signal at the center of each individual centriole and lines were rotated at 45° intervals (eight measurements per centriole). The mean of all line scans is plotted along with the standard deviation. The slope of these lines is plotted (Y axis on right) and the vertical lines indicate where the fluorescence intensity begins to decrease in each condition. E) Microtubules emanate primarily from the mother centriole when SON is depleted. Tomographic slices with modelled centrioles, procentrioles and microtubules overlayed. F) Distances of the microtubule minus ends to the center of the centriole. N = siControl, 340; siSON, 379 from four and five tomographic models, respectively. G) Katanin localization around the mother centriole is dependent upon SON. SIM images of siRNA treated S phase arrested cells induced for PLK4 expression stained for Katanin p80 (KATNB1) (red). H) Fluorescence quantification of Katanin p80 (KATNB1) in a 1 μm radius circle around the centrosome from widefield images. N = siControl: 184 centrosomes; siSON 177 centrosomes from three biological replicates. Scale bars = 1 μm for A), C) and G). Scale bars = 200 nm in E). p values are less than 0.0001. Bar and error bars: mean and standard deviation.

We next analyzed the relationship between microtubule minus ends and centrioles in EM tomograms. Consistent with the fluorescent image analysis, microtubules on average originated closer to the mother centriole upon SON depletion (408 nm) as compared to control conditions (483 nm) (Figure 7E, F, Figure Supplement 7A, B). Although centriolar satellites are difficult to discern without a protein marker (Kubo et al., 1999), the cytoplasmic content in SON depleted cells is markedly different from that of control cells with a dramatic increase in the number of vesicles present near the centrosome (See Movie S2, Movies S4-S7). The increase in vesicles in this region may be due to the absence of centriolar satellites which can no longer preclude vesicle encroachment near the centrosomes.

To explore how SON influences the microtubule network organized by centrosomes, we examined the localization of the p80 subunit of the Katanin complex responsible for its centrosomal localization (McNally et al., 2000). Katanin p80 is encoded by *KATNB1* whose mRNA splicing is affected by SON (Figure Supplement 4C) (Ahn et al., 2011). Furthermore, the absence of Katanin p80 stabilizes microtubules near the centrosome (Hu et al., 2014). *KATNB1* mRNA abundance is reduced in siSON to 18% of the siControl samples, and splicing efficiency is also reduced, particularly at the 3’ end of the gene (Figure Supplement 4C). Katanin p80 localization is enriched around the centrosome in the region between the mother centriole and procentrioles. This localization is reduced when SON is depleted (Figure 7G, H). We suggest that microtubules are actively remodeled by centrosome localized Katanin p80 to ensure a transition from the mother centriole to nascent procentrioles. When SON is depleted, Katanin is removed from the centrosome thereby increasing microtubule nucleation directly from the mother centriole. Together, with the decreases in centriolar satellites and Centrobin, procentriole elongation and microtubule triplet formation are inhibited.

## Discussion

We have shown that the function of the splicing factor SON is required for centriole assembly, independent of its effects on the cell cycle. Although centriole assembly initiates in the absence of SON, procentrioles do not form completely. Correct splicing of the gene encoding the centriole assembly factor Centrobin, which is required for centriole elongation, is dependent on SON activity. The absence of Centrobin does not explain the absence of Centrin incorporation (a marker of more mature procentrioles) observed with SON depletion but may contribute to the lack of triplet microtubules in the procentrioles. SON is also responsible for promoting proper mRNA splicing of *CEP131* which encodes a core centriolar satellite protein. SON depletion profoundly disrupts the distribution of centriolar satellites and Pericentrin trafficking granules around the centrosome. Yet even the depletion of CEP131 and Pericentrin alone or in combination cannot fully recapitulate the centriole assembly defect caused by loss of SON. SON also affects the splicing of *KATNB1*, which encodes for the p80 centrosomal targeting subunit of katanin. KATNB1 changes at the centrosome may explain the increase in microtubule density near centrosomes in SON depletion cells. Together, this data suggests that SON has an important role in controlling factors critical for trafficking particles around the centrosome and the microtubule cytoskeletal network upon which they rely.

Whether centriolar satellites are required for centriole assembly appears to be cell type dependent as different centriole assembly responses to centriolar satellite disruption are observed. Cell lines developed from cancer cells (HeLa, U2OS) utilize centriolar satellite transport for centriole assembly, whereas hTERT immortalized cells (RPE-1) forgo this pathway (Hori et al., 2015; Kodani et al., 2015). In this study, using RPE-1 cells, we observed close association of centriolar satellites with assembling procentrioles, but depletion of satellites (through CEP131 depletion) and Pericentrin associated trafficking particles caused reductions in centriole assembly, but did not eliminate it. The resilience in centriole assembly observed in this study could be due to the S phase arrest and PLK4 induction used in these experiments, which may make the process unnaturally robust. Alternatively, it could reflect the lack of reliance on centriolar satellites for centriole assembly in RPE-1 cells.

There is, however, increasingly strong evidence that centriolar satellites are required for primary cilia formation and function (Chamling et al., 2014; Hori et al., 2016; Kurtulmus et al., 2015; Odabasi et al., 2019). Given that SON has such a profound effect on centriolar satellites, it is likely that defects in SON are also deleterious to cilia. This is consistent with the discovery that *de novo* mutations in *SON* are associated with intellectual disabilities (J.-H. Kim et al., 2016; Tokita et al., 2016), which can be caused by ciliary defects (Youn & Han, 2018). *SON* resides on human chromosome 21 and is therefore one of the genes whose dosage is altered in Down Syndrome. Given that individuals with Down Syndrome have defects in their cilia and cilia-dependent signaling (Galati et al., 2018; Moldrich et al., 2007; Roper et al., 2006), the role that *SON* dosage plays in cilia formation and function requires investigation. SON’s activities could be required constitutively or modulated for unique cell cycle and developmental programs. Our data indicate that SON can effectively control centrosome assembly, which would have consequences for both the cell cycle, and development. SON was identified to be important for maintaining human embryonic stem cell (hESC) identity (Chia et al., 2010), and its depletion led to hESC differentiation into a fibroblast-like state with a consequent downregulation of pluripotency-associated genes, upregulation of differentiation-associated genes and no change to housekeeping genes (Lu et al., 2013). As pluripotency requires high fidelity cell divisions and centrosomes, SON control of centrosome assembly could contribute to stemness and provide a powerful avenue to limit both cell division during differentiation and centrosomal function as a dominant MTOC in cell types that utilize non-centrosomal MTOCs in their differentiated state.

In conclusion, SON activity has a strong positive impact on trafficking particles around the centrosome, including centriolar satellites and cytoplasmic Pericentrin granules, that play an important role in completing procentriole assembly. SON is required for the correct mRNA splicing of *CEP131* and the efficient splicing of *PCNT*. SON is also involved in the remodeling of the underlying microtubule network upon which centriolar satellites and Pericentrin trafficking particles rely, perhaps through its involvement in the correct splicing of *KATNB1*, the centrosomal targeting subunit of the katanin complex. In addition, SON is important for proper splicing of *CNTROB*, the gene encoding Centrobin which is important for centriole elongation and microtubule triplet formation. Together, SON acts as a critical nexus for a powerful, multifactorial control of centriole assembly through its influence on genes required for the microtubule network and centriolar satellites that support centriole assembly.

## Data Availability

Raw and processed RNA sequencing data for this study have been deposited at NCBI GEO under accession GSE164278.

## Methods

### Cell culture growth conditions, small interfering RNA treatments and DNA transfections

RPE-1/Tet-PLK4 GFP-CETN (the kind gift of M.B. Tsou) (Hatch et al., 2010) were grown in DMEM/high glucose with Glutamine (Cytiva or Gibco) with 10% tetracycline free fetal bovine serum (Peak Serum) supplemented with penicillin and streptavidin (Gibco) at 37°C with 5% CO_2_. For PLK4 induction experiments, cells were plated onto 12 mm circular cover glass, No. 1.5 (Electron Microscopy Sciences) coated with collagen (Sigma, C9791) in 24-well plates at 11,000 to 12,500 cells per well. The following day cells were treated with small interfering RNAs using the Lipofectamine RNAiMAX Transfection Reagent (ThermoFisher Scientific) following the manufacturer’s instructions. Transfection complexes were removed from cells after 5 hours. 24 hours after transfection complexes were added, cells were arrested with 1.6 µg/mL aphidicolin (Cayman Chemical Company). PLK4 was induced with 1 µg/mL Doxycycline (Sigma-Aldrich) seven hours after aphidicolin arrest was initiated. Cells were fixed seventeen hours after PLK4 induction.

siRNAs utilized were SON: Stealth P5A8 (all experiments), GCGCUCUAUGAUGUCAGCUUAUGAA; Stealth P5A7, CCGAUCUAUGAUGUCAUCUUAUACU; CEP131: Stealth HSS146116, CAGAGUGCCAGGAAUGCGGCAGCCU; CNTROB: Silencer Select s42058 GCAUUGGAUUCAGAGCAUAtt; Stealth siRNAs were used at 50 nM, and Silencer Select siRNAs were used at 10 nM (Thermo Fisher Scientific). Pericentrin was targeted with an siGenome Smartpool (Dharmacon) at 5 nM.

DNA transfections were done with Lipofectamine 2000 (Thermo Fisher Scientific) according to the manufacturer’s instructions.

### Immunofluorescence

Cells were fixed in cold MeOH for eight minutes, washed three times in PBS, and blocked in Knudsen buffer (1× PBS, 0.5% bovine serum albumin, 0.5% NP-40, 1 mM MgCl2, 1 mM NaN3) for 1 hour. Cells stained for tubulin were fixed with Glutaraldehyde and Formaldehyde as previously described (Canman et al., 2000). Antibody staining was conducted at room temperature for 1 to 2 hours. Samples were washed 3 times for 5 minutes in PBS prior to secondary antibody and DNA staining in Knudson buffer for 1 hour, except for samples stained for tubulin in which fixes, staining and washes were conducted in PHEM buffer. Primary antibodies: 1:2000 α-γ-tubulin (Sigma, DQ19); 1:200 α-FLAG (Sigma, M2); 1:2500 α-CEP152 (Bethyl A302-480A); 1:2000 α-CEP192 (the generous gift of A. Holland, Department of Molecular Biology and Genetics, Johns Hopkins University School of Medicine, Baltimore, MD); 1:2500 α-STIL (Bethyl SIL A302-441A); 1:2000 α-SAS-6 (Bethyl A301-802A); 1:500 α-CPAP (Proteintech CENPJ 11517-1-AP); 1:300 α-CEP135 (Proteintech 24428-1-AP); 1:4000 α-SON (Thermo Fisher Scientific PA5-65107); 1:2000 α-Pericentrin (Abcam ab4448); 1:1000 α-Centrobin (Proteintech 26880-1-AP); 1:10,000 α-CEP131 (the generous gift of J. Reiter, Department of Biochemistry and Biophysics, University of California, San Francisco School of Medicine, San Francisco, CA); 1:2000 α-PCM1 (Bethyl A301-150A) α-α-tubulin (Sigma DM1A); 1:50 α-KATNB1 (Proteintech 14969-1-AP). Secondary staining: 1:1000 Alexa anti-Rabbit 594, Alexa anti-Mouse 594, Alexa anti-Guinea Pig 647, and Hoechst 33342 (Thermo Fisher Scientific). Coverslips were mounted on Citifluor (Ted Pella) and sealed with clear nail polish.

### Fluorescence Imaging

Images were collected using a Nikon TiE (Nikon Instruments) inverted microscope stand equipped with a 100× PlanApo DIC, NA 1.4 objective. Images were captured using an Andor iXon EMCCD 888E camera with 0.3μm Z steps. Structured Illumination Microscopy images were acquired using a Nikon SIM (N-SIM) with a Nikon Ti2 (Nikon Instruments; LU-N3-SIM) microscope equipped with a 100× SR Apo TIRF, NA 1.49 objective. Images were captured using a Hamamatsu ORCA-Flash 4.0 Digital CMOS camera (C13440) with 0.1μm Z step sizes. Reconstructions were generated with Nikon Elements. All images were collected at 25°C using NIS Elements software (Nikon). Raw SIM images were reconstructed using the image slice reconstruction algorithm (NIS Elements).

### Centriole counts

GFP**-**Cetn fluorescence was utilized to count centrioles manually using the 100× PlanApo DIC, NA 1.4 objective with the optivar in on the Nikon TiE inverted microscope stand, providing 1.5X magnification.

### Image analysis

Image analysis was done using Fiji (Schindelin et al., 2012). Image stacks were maximum intensity projected. Fluorescence intensities of centrosomes were measured within a 1 μm radius circle encompassing each individual centrosome (mother and daughter centrioles). A measurement of in cell intensity near each centrosome was taken for subtraction of non-centrosomal signal. To measure the fluorescence intensity of proteins with satellite and trafficking populations, fluorescence was measured within a 5 μm radius circle encompassing both centrosomes and the surrounding cytoplasm. A similar circle was measured within the cell that did not include the centrosomes to calculate the in-cell background. If subtraction of background signal generated a negative result for fluorescence, it was converted to 0.

For radial fluorescence intensity analyses, we utilized the algorithm described in Sankaran et al (Sankaran et al., 2020) using the PCM or microtubule analysis tool. Briefly, maximum intensity projected images were selected for analysis. Centrosomes and cells identified by the algorithm were manually checked for accuracy prior to the algorithm calculating the in-cell radial fluorescence intensity.

SON total nuclear fluorescence intensity was measured in Fiji by subtracting the average background (calculated from all images analyzed in a 5μm radius circle outside of the cells) from each image. The DNA signal (Hoechst 33342) was used to define the nuclear regions using the threshold tool and converted to a binary image. The erode and dilate binary functions were applied once to smooth the regions prior to applying the create selection function. The selection was added to the region of interest manager and then split into its separate constituents. Only regions that encompassed one full nucleus were retained. These regions were transferred to the SON channel where the fluorescence intensity was measured.

To assess total fluorescence within a cell, background was subtracted as described in the previous paragraph. The Cetn channel was thresholded to generate a binary image encompassing all the cell boundaries. This was subjected to the erode, dilate and fill holes binary functions followed by applying the create selection function. The selection was added to the region of interest manager and then transferred to the channel to be quantified.

To measure the proximity of microtubules to centrosomes in structured illumination microscopy reconstructions, line scan measurements of fluorescence intensity were employed beginning at the center of the centriole signal. The line was rotated 45°, generating eight measurements for each centriole. Each line scan measurement was then normalized to one at the initiating measurement at the centriole center prior to averaging all the line scans for each condition.

### Flow cytometry

Cells were grown, treated with si RNAs and S phase arrested as described above in 6 well plates (inoculum of 64,000 cells/well). To collect cells, media, PBS wash and trypsin dissociated cells were combined into one tube per condition, spun and washed with PBS and resuspended in Krishan stain (3.8mM Sodium citrate, 69nM Propidium iodide, 0.01% NP40, 0.01mg/mL RNaseA) (Krishan, 1975) and incubated at 4°C until run and analyzed by the University of Colorado Cancer Center Flow Cytometry Shared Resource on a Beckman Coulter FC500 flow cytometer. Cell cycle analysis was performed using ModFit LT software (Verity Software House, Topsham, ME).

### RNA isolation, RT-PCR, and library preparation

RNA was isolated using the manufacturers recommended protocols in three ways for the three replicates averaged in Figure 4C. Cytoplasmic RNA was isolated using the RNeasy Minikit (Qiagen) and the supplementary protocol provided by the manufacturer that includes a cell lysis step using buffer RLN and a step to separate nuclei from the cytoplasm by gentle centrifugation. RNA was also isolated using the Monarch Total RNA isolation miniprep kit (New England Biolabs). RNA was also extracted using the TRI Reagent (Thermo Fisher Scientific). cDNA was generated using 1 μg RNA, random hexamers (Thermo Fisher Scientific SO142) and SuperScript III Reverse Transcriptase (Thermo Fisher Scientific 18080044). Primers for PCR were ACTB-F: AGAGCTACGAGCTGCCTGAC; ACTB-R: AGCACTGTGTTGGCGTACAG; CNTROB-4F: TGCAAGACTTGTCTCCATCTAGCTC; CNTROB-5R: TTGTCCAGTTGTTCAATCATGGTATCTTTC; CNTROB-9F: AGAAGAGCCAGAGGGAAGCC; CNTROB-10R: TTGCCGTAGGCTGCTCTCC; CEP131-4F: ACGGAGCCCACAGACTTCC; CEP131-6R: CGCAGTTGCCCACTGCTC; CEP131-14F: TGGGGTCCGAGGTGAGC; CEP131-15R: GCTGGATGGTGGCCTCG. Bands were quantified using Fiji (Schindelin et al., 2012).

RNA used for polyA selected library generation for mRNA sequencing was isolated with the Monarch Total RNA isolation miniprep kit (New England Biolabs). Libraries were constructed using the Nugen Universal Plus mRNA-SEQ library construction kit (Nugen 0508) and sequenced on an Illumina NovaSEQ 6000 sequencer by the Genomics and Microarray shared resource at the University of Colorado Cancer Center.

### Bioinformatics

RNA-seq libraries were sequenced on an Illumina NovaSeq 6000 (2×150) to a depth of 60-100 million paired-end reads. Reads were trimmed using cutadapt (v1.16) to remove adapters and aligned to the hg38 genome using STAR (v2.5.2a, (Dobin et al., 2013)). Gene counts were calculated using featureCounts (v1.6.2, (Liao et al., 2014)). Differentially expressed genes were identified for cells expressing SON siRNA and siControl cells (three replicates each) using DESeq2 (v1.28.1, (Love et al., 2014)). To compare annotated gene ontology terms alongside our custom gene lists, we generated a custom Gene Matrix Transposed (.gmt) file containing cellular component gene ontology terms (v7.1) along with the Centrosome Database and centriolar satellites gene lists. To identify GO terms, genes downregulated after SON knockdown (adjusted p-value < 0.05, log2 fold change <-0.4) were sorted by p-value and submitted as an ordered query using the R package gprofiler2 (v0.2.0, (Kolberg et al., 2020)).

Genes with changes in alternative splicing events were identified using rMATS (v4.1.0, (Shen et al., 2014)). Splicing events were identified using both junction and exon read counts and filtered to only include those with an FDR <0.05 and where the absolute inclusion difference was >0.4. GO terms were identified as described above.

Changes in alternative splicing were also analyzed using the software package, MAJIQ (v1.1.7a, (Vaquero-Garcia et al., 2016)). MAJIQ calculates the change in PSI (percent spliced in) for splicing events between samples. The MAJIQ results were filtered to only include those where the change in PSI after SON knockdown was >0.2. GO terms were identified as described above.

### Electron microscopy tomography

Cells were grown on sapphire discs and prepared for electron microscopy using high pressure freezing and freeze substitution (McDonald et al., 2010). Briefly, 3mm sapphire discs (Technotrade International) were coated with gold. A large F was then scratched into the surface to help orient the cell side. The disks were coated with collagen, sterilized under UV light, and cells were plated for culturing as described above. Monolayers of cells grown on sapphire discs were then frozen using a Wohlwend Compact02 high pressure freezer (Technotrade International). The frozen cells were then freeze substituted in 1% OsO4 and 0.1% uranyl acetate in acetone at −80°C for 3 days then gradually warmed to room temperature. The discs were then flat embedded in a thin layer of Epon resin and polymerized at 60°C. Regions containing cells were identified in the light microscope, and a small square of resin containing the cells was excised and remounted onto a blank resin block. The cells were then sectioned en face and serial, thick sections (250-300nm) were collected onto formvar-coated slot grids. Some grids were post stained with 2% uranyl acetate and Reynold’s lead citrate. 15 nm colloidal gold (BBI International) was affixed to the section surface to serve as alignment markers.

Tomography was performed using a Tecnai F30 microscope operating at 300 kV (Thermo Fisher Scientific, Waltham, MA). Dual axis tilt series were collected over a +/− 60° range using the SerialEM image acquisition software (David N. Mastronarde, 2005) and a Gatan OneView camera (Gatan, Inc., Pleasanton, CA). For some data sets, tilt series were collected from 2-4 serial sections to reconstruct a larger volume of cell data. Tomograms were computed, serial tomograms joined and cellular features were modeled using the IMOD 4.9 software package (https://bio3d.colorado.edu/imod/; (Kremer et al., 1996; D. N. Mastronarde, 1997)).

MTs, centrioles, procentrioles and positions of MT ends were manually traced in these reconstructions using the 3dmod program in the IMOD software package (Kremer et al., 1996). The slicer window in 3dmod was used to rotate a slice of image data containing the procentriole to view the microtubule organization in cross or longitudinal view. Models were projected in 3D to show the arrangement of the forming procentrioles in the centrosomes and the position of microtubules and their ends within the volume. The places of close approach between microtubule ends to the centriole surface were identified with the MTK program in the IMOD suite. For this analysis, a single point at the minus end of each MT was modeled. The mother centriole was modeled as a series of closed contours to represent its surface. The MTK program uses models of microtubule ends to identify points of close apposition to the centriole and outputs a model object at each point of close approach. Measurements of the lengths of the close approach model contours were then extracted using the imodinfo program in the IMOD suite. The 3D distance between MT ends and a single reference point marking the center of the centriole was determined using the imod-dist program in the IMOD suite. The imod-dist program was then used to compute the 3D distances between the centriole and the locations of microtubule ends. In total, centrosomes from 4 siControl and 4 siSON cells were reconstructed and modeled in a total of 10 tomograms.

### Plasmid construction

CEP131 first strand cDNA was isolated using the BamHI-CEP131-3’ primer (AACGGATCCTCACTTGGTACTTGGCGTGG) and SuperScript III reverse transcriptase (Thermo Fisher Scientific), followed by PCR amplification with EcoRI-CEP131-5’ (ACCGAGAATTCCATGAAAGGCACCCGGGC) using KAPA HiFi DNA polymerase (Roche). The product was digested with EcoRI and BamHI and ligated into similarly digested pEGFP-C1 (Clontech). The resulting plasmid was then digested with NheI and XhoI to remove GFP and replace with mCherry from similarly digested pmCherry-C1 (Clontech).

## Acknowledgments

We thank Pierre Gönczy and Fernando Balestra for reagents and discussions, Andrew Holland and Jeremy Reiter for antibodies and Meng-Fu Bryan Tsou for the RPE1/Tet-PLK4/GFP-CETN cell line. Electron microscopy was done at the University of Colorado, Boulder EM Services Core Facility in the MCDB Department, with Garry Morgan providing specimen preparation. We also thank Marisa Ruehle for critical readings of the manuscript and the Pearson lab for discussions. J. T. M. was supported by the Boettcher Foundation. C.G.P. is funded by the American Cancer Society (RSG-16-157-01-CCG) and the National Institute of General Medical Sciences (R01GM099820 and R01GM132132).

## Supporting Material

**Figure Supplement 1.**
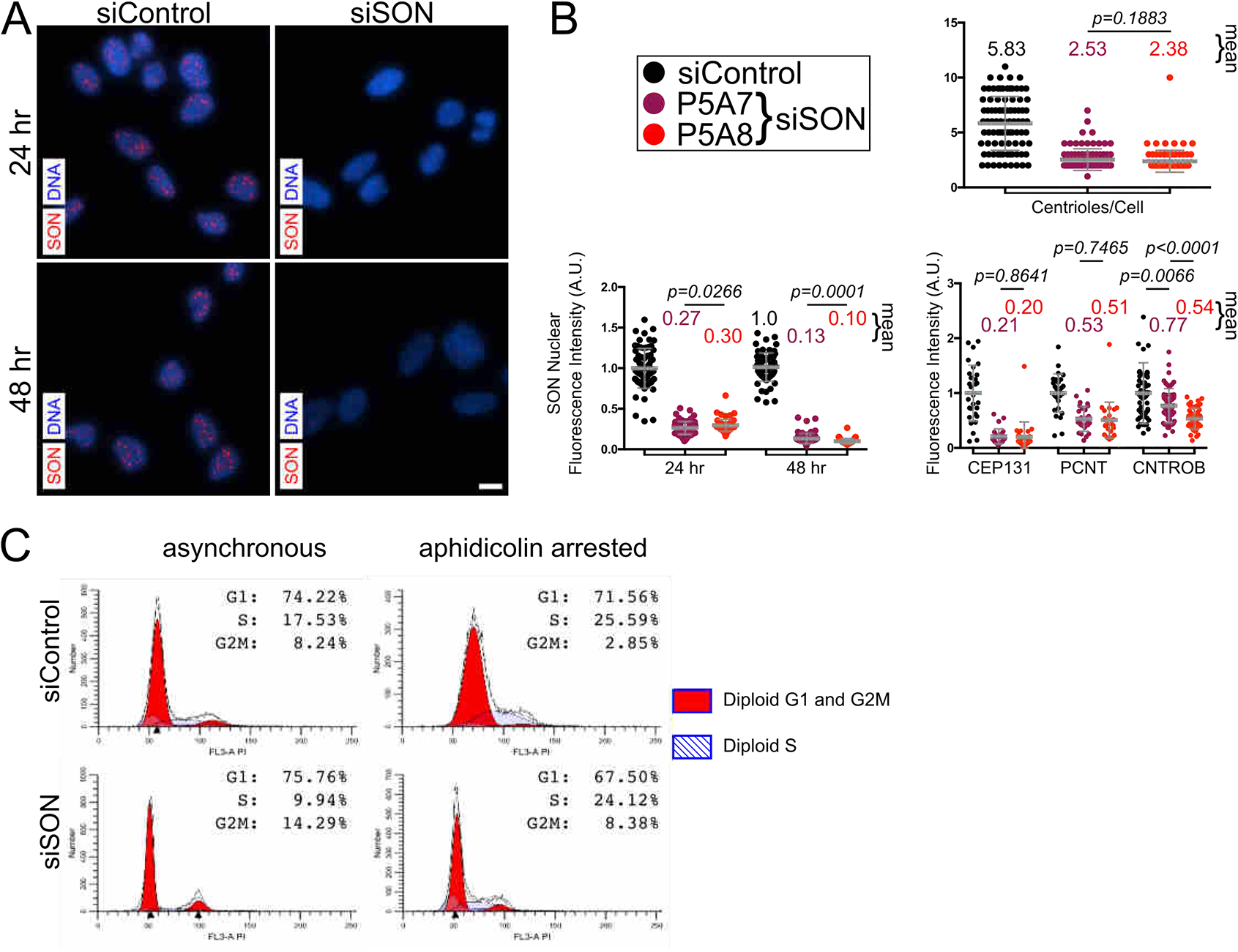
Two SON siRNA treatments are similar and S phase arrest is similar to siControl treated cells. A) Cells stained for SON (red) and DNA (Hoechst stain, blue) treated with either control or SON siRNAs (P5A8) for 24 or 48 hours. Scale bar = 10 μm. B) Comparisons of two SON siRNAs, including quantification of the SON nuclear signal (N = 24hr: siControl, 71; siSON P5A7, 71 cells; siSON P5A8, 72 cells; 48hr: siControl 72 cells; siSON P5A7, 62 cells, siSON P5A8, 64 cells), centrioles per cell (N = 90 cells per condition), CEP131 and Pericentrin fluorescence intensities within a 5 µm radius circle (N = siControl, 29; siSON P5A7, 28; siSON P5A8, 27 cells) and the Centrobin fluorescence intensities within a 1 µm radius circle (N = siControl, 56; siSON P5A7, 60; siSON P5A8, 54 centrosomes). Bar and errror bars: mean and standard deviation. Data is from one biological replicate. p values are < 0.0001 unless otherwise notated. C) Flow cytometric analysis of DNA content of cells with or without aphidicolin S phase arrest. Percentages of cells in different cell cycle stages are indicated.

**Figure Supplement 2.**
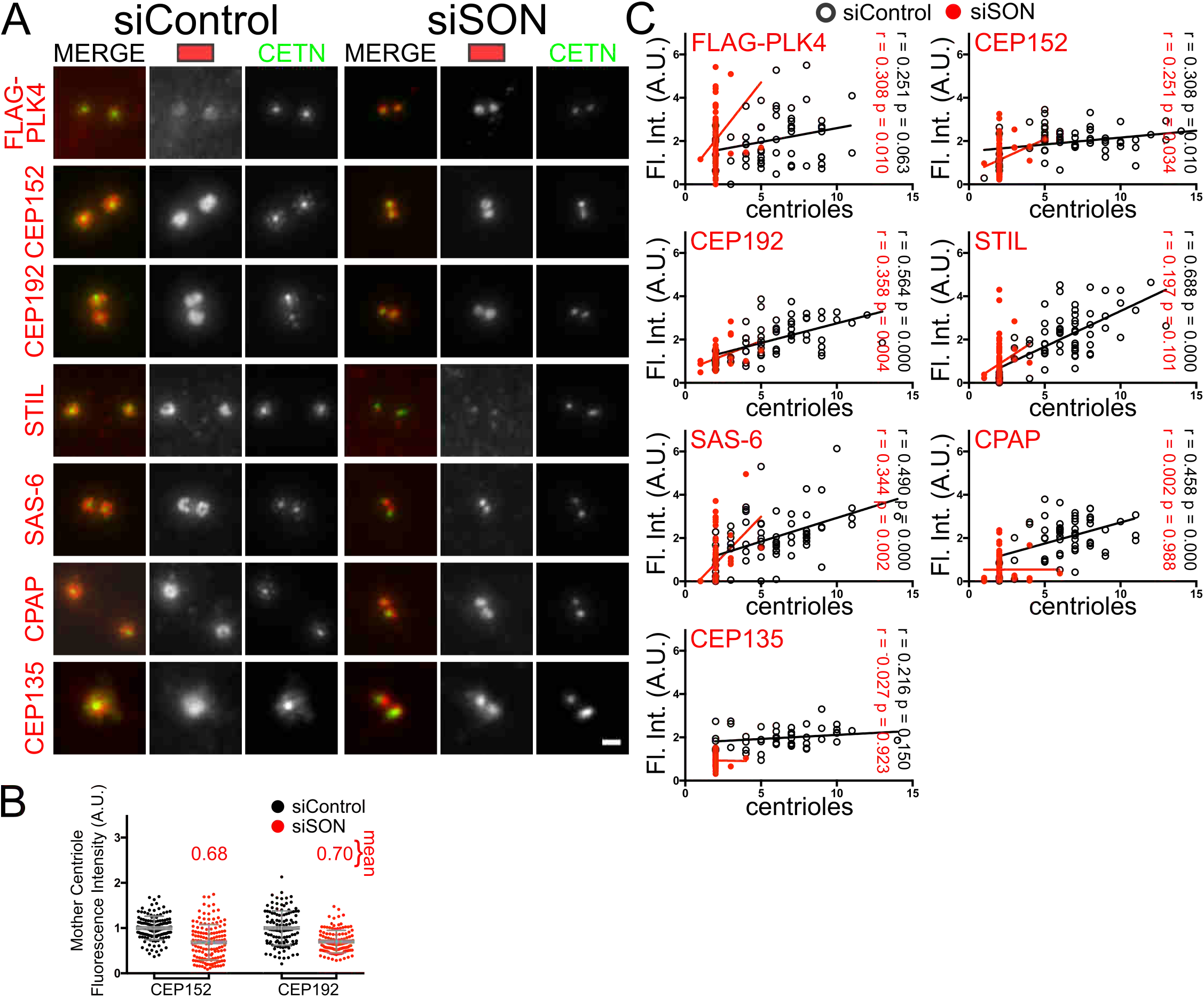
SON depletion reduces the amount of early centriole assembly factors. A) Representative widefield images used for quantification of the indicated early centriole assembly factors. Scale bar = 1 μm. B) Fluorescence quantification of CEP152 and CEP192 at the mother centriole (0.39 μm square). N = Cep152: siControl, 134 centrioles; siSON, 140 centrioles; Cep192: siControl, 122 centrioles; siSON, 121 centrioles from two biological replicates. C) Correlated plots of the number of centrioles and the fluorescence intensity within an individual cell, stained for the indicated core centriole assembly factors. siControl = black circles, siSON = red dots. Linear regressions are plotted. r and p values are indicated in the appropriate colors. N = FLAG-PLK4: siControl, 56 cells; siSON, 69 cells; Cep152: siControl, 70 cells; siSON, 72 cells; Cep192: siControl, 62 cells; siSON, 63 cells; STIL: siControl, 68 cells; siSON, 70 cells; Sas-6: siControl, 67 cells; siSON, 77 cells; CPAP: siControl, 60 cells; siSON, 64 cells; Cep135: siControl, 46 cells; siSON, 60 cells. Data is compiled from two biological replicates.

**Figure Supplement 3.**
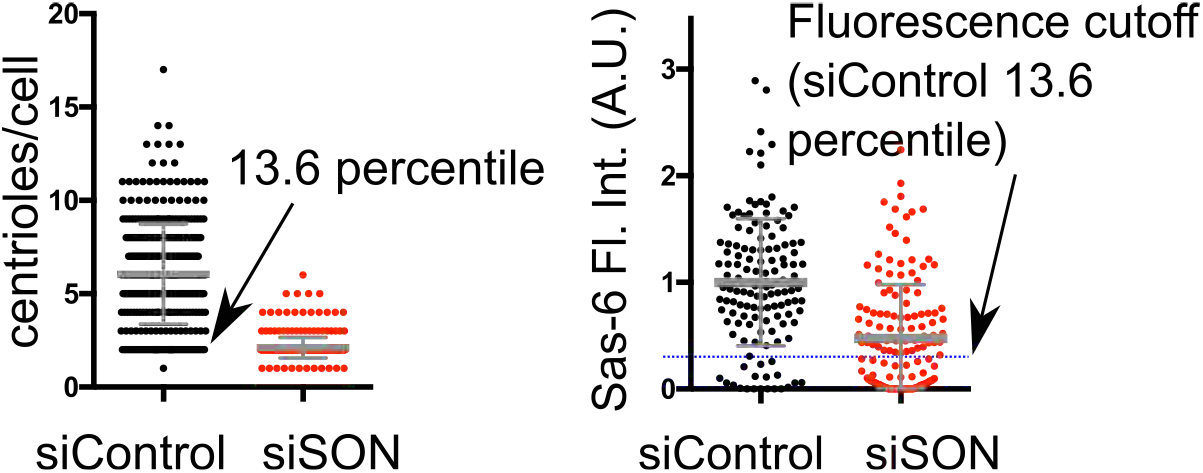
SON depletion is permissive for early core centriole assembly events. Determination of a SAS-6 fluorescence intensity cutoff to estimate the initiation of new centriole assembly. Centrin counts were determined in C) (n = 383 siControl cells) and SAS-6 fluorescence is also shown in Figureure 2B.

**Figure Supplement 4.**
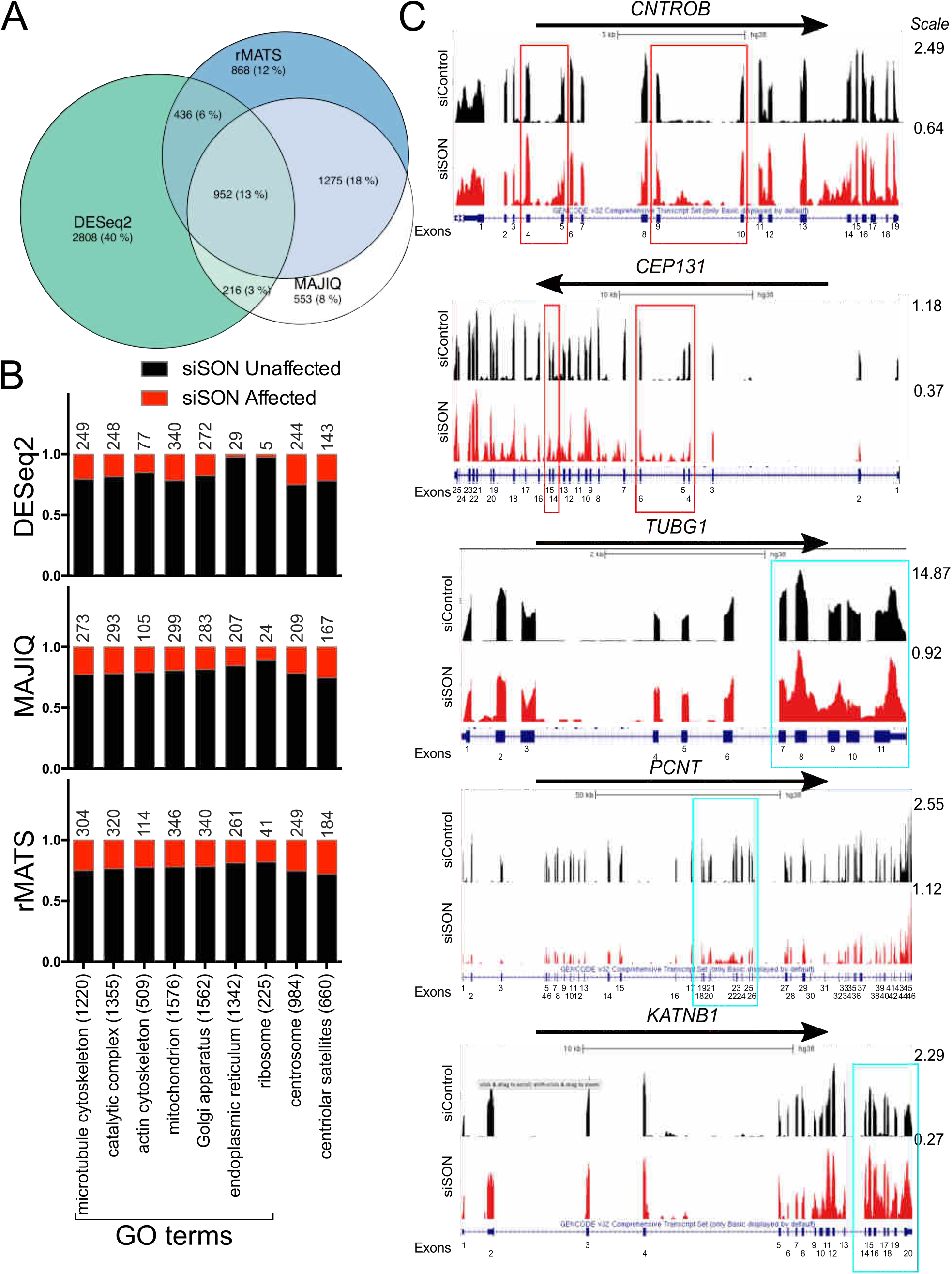
SON depletion affects the splicing of centrosomal, centriolar satellite and microtubule cytoskeletal mRNAs. A) A Venn diagram displaying the overlap of genes with reduced expression and alternative splicing when SON is depleted as determined by DNASeq2, MAJIQ and rMATS. B) Fraction of genes with reduced expression or alternatively spliced in selected GO terms that encompass major cellular structures when SON is depleted. The centrosome database and centriolar satellite proximity list are also included (centrosome and centriolar satellites). The term size is listed at the bottom, and the number of genes affected by SON are above each bar. C) Counts from RNA sequencing from selected genes affected by SON depletion. Arrows indicate the direction of the transcript relative to the genome. Blue boxes indicate exons, which are numbered beneath. Red boxes indicate regions which were investigated by reverse transcription PCR in Figureure 4. Cyan boxes indicate regions of altered splicing. Scales are indicated on the righthand side and unique to each plot.

**Figure Supplement 5.**
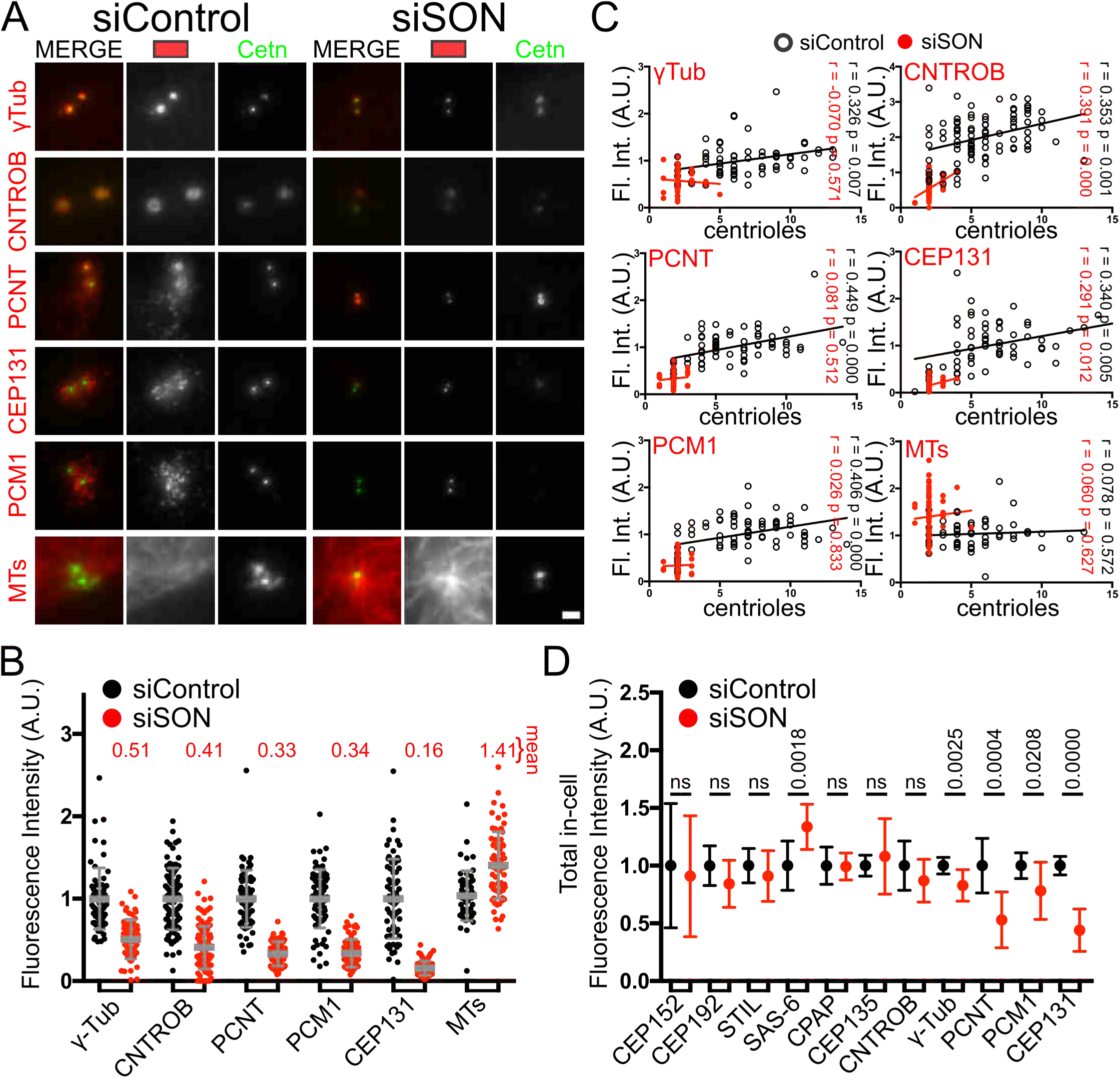
SON splicing targets affect centriolar satellites and trafficking granules. A) Representative widefield images used for fluorescence quantification of the indicated proteins (red). Scale bar = 1 μm. B) Fluorescence quantification of the indicated proteins in a 5 μm radius circle around the centrosome. N = γ-Tubulin: siControl, 65 cells; siSON, 63 cells; Centrobin (CNTROB): siControl, 90 cells; siSON 100 cells; Pericentrin (PCNT): siControl, 65 centrosomes; siSON, 66 centrosomes; PCM1: siControl, 74 centrosomes; siSON, 67 centrosomes; Cep131: siControl, 66 centrosomes; siSON, 73 centrosomes; microtubules (MTs): siControl, 55 centrosomes; siSON 68 centrosomes. Bar and error bars: mean and standard deviation. Data compiled from two biological replicates. C) Correlated plots of the number of centrioles and the fluorescence intensity within an individual cell, stained for the indicated marker. siControl = black circles, siSON = red dots. Linear regressions are plotted. r and p values are indicated in the appropriate colors. N = γ-Tubulin: siControl, 67 cells; siSON, 68 cells; Centrobin (CNTROB): siControl, 44 cells; siSON, 56 cells; Pericentrin (PCNT): siControl, 65 cells; siSON, 66 cells; CEP131: siControl, 66 cells; siSON, 73 cells; PCM1: siControl, 74 cells; siSON, 67 cells; microtubules (MTs): siControl, 55 cells; siSON, 68 cells. Data is compiled from two biological replicates. D) Total in cell fluorescence intensities. Fluorescence was measured simultaneously for all cells in each image, and 10 images total were used for each marker except for CEP131 where 9 images were used and Centrobin (CNTROB), where 16 images were used. Data is compiled from two biological replicates. Bars and error bars: mean and standard deviation. p < 0.0001 unless otherwise indicated. ns = not significant.

**Figure Supplement 6.**
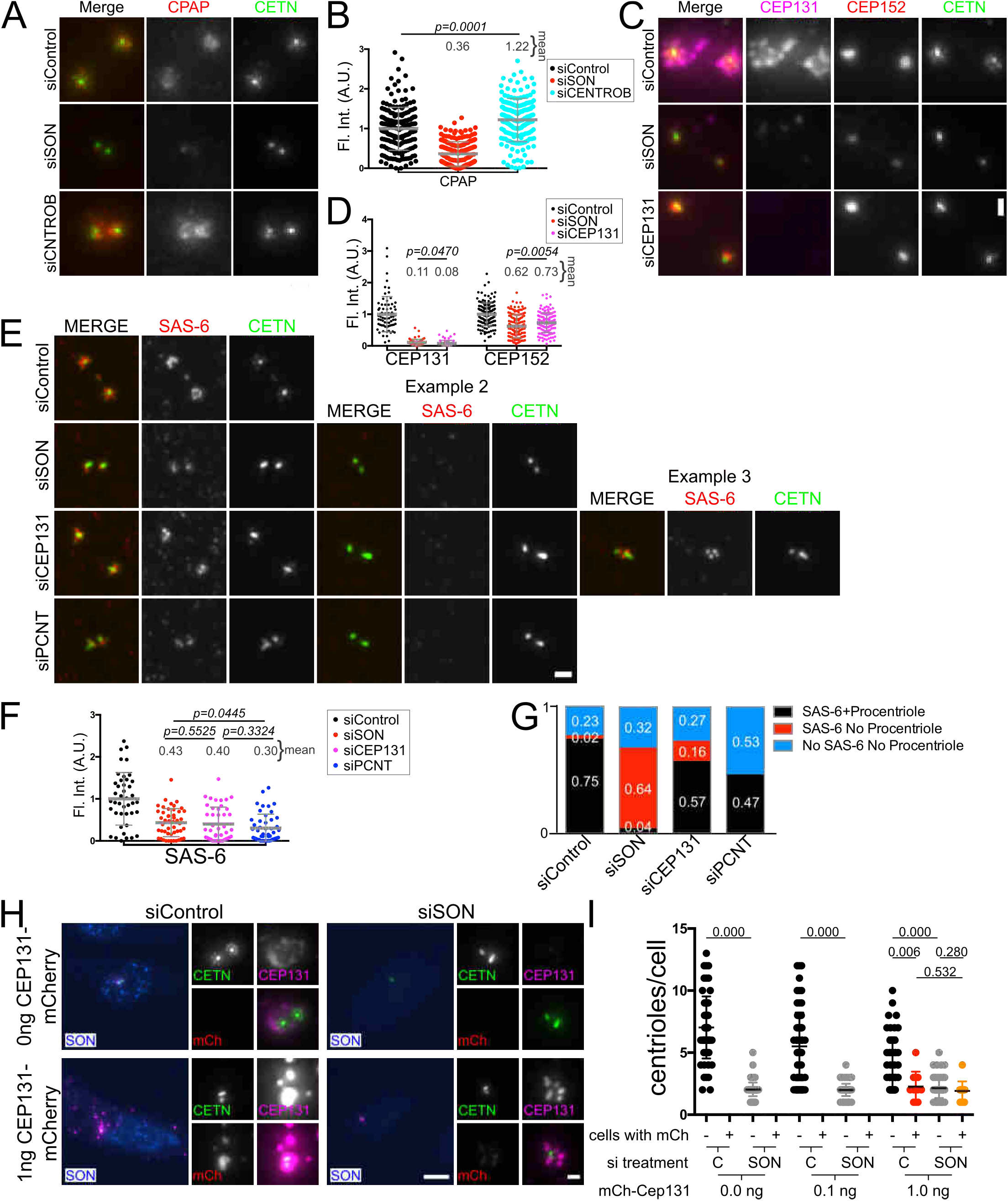
Depletion of SON splicing targets partially phenocopy SON depletion. A) Centrobin depletion does not reduce CPAP fluorescence intensity at the centrosome. Widefield images of cells treated with control, *SON*, or Centrobin (*CNTROB*) siRNAs and stained for CPAP. B) Quantification of CPAP fluorescence intensity within a 1 μm radius circle around the centrosome. N = siControl: 181 centrosomes; siSON: 196 centrosomes; siCNTROB: 185 centrosomes. Bar and error bars: mean and standard deviation. C) CEP131 depletion reduces CEP152 at the centrosome. Widefield images of cells treated with control, SON, or CEP131 siRNAs and stained for CEP131 and CEP152. D) Quantification of CEP131 and CEP152 fluorescence intensities within a 5 μm radius circle or 1 μm radius circle respectively. N = siControl: CEP131, 69 cells; CEP152, 138 centrosomes; siSON: CEP131, 67 cells; CEP152, 150 centrosomes; siCEP131: CEP131, 64 cells; CEP152, 137 centrosomes. Compiled from two biological replicates. Bar and error bars: mean and standard deviation. E) CEP131 and SON depletion can delay completion of procentriole assembly after initiation of procentriole assembly. SIM images of cells treated with control, *SON*, *CEP131* or Pericentrin (*PCNT*) siRNAs and stained for SAS-6. F) Quantification of SAS-6 fluorescence intensity within a 1 μm radius circle around the centrosome. N = siControl: 42 centrosomes; siSON: 48 centrosomes; siCEP131: 44 centrosomes; siPCNT: 47 centrosomes from one biological replicate. G) 75 cells from Control, *SON*, *CEP131* or *PCNT* siRNA treated cells were classified based on the presence of SAS-6 and procentrioles. H) Widefield fluorescence microscopy images of siControl or siSON cells that either went through a transfection with no DNA or 1 ng of mCherry-CEP131 DNA. SON antibody = blue, GFP-Cetn autofluorescence = green, mCherry=Cep131 autofluorescence = red, Cep131 antibody = magenta. Scale bar for whole cells = 10 μm and for insets = 1 μm. I) Quantification of centrioles per cell in siControl and siSON cells after transfection with the indicated amount of mCherry-CEP131 DNA. mCherry signal was used to identify cells transfected with the mCherry-CEP131 construct. N = siControl: 0 ng DNA, 62 cells; 0.1 ng DNA untransfected, 76 cells; transfected, 0 cells; 1 ng DNA untransfected, 52 cells; transfected, 11 cells; siSON: 0 ng DNA, 70 cells; 0.1 ng DNA untransfected, 76 cells; transfected, 0 cells; 1 ng DNA untransfected, 57 cells; transfected, 13 cells. Bars and error bars: mean and standard deviation.

**Figure Supplement 7.**
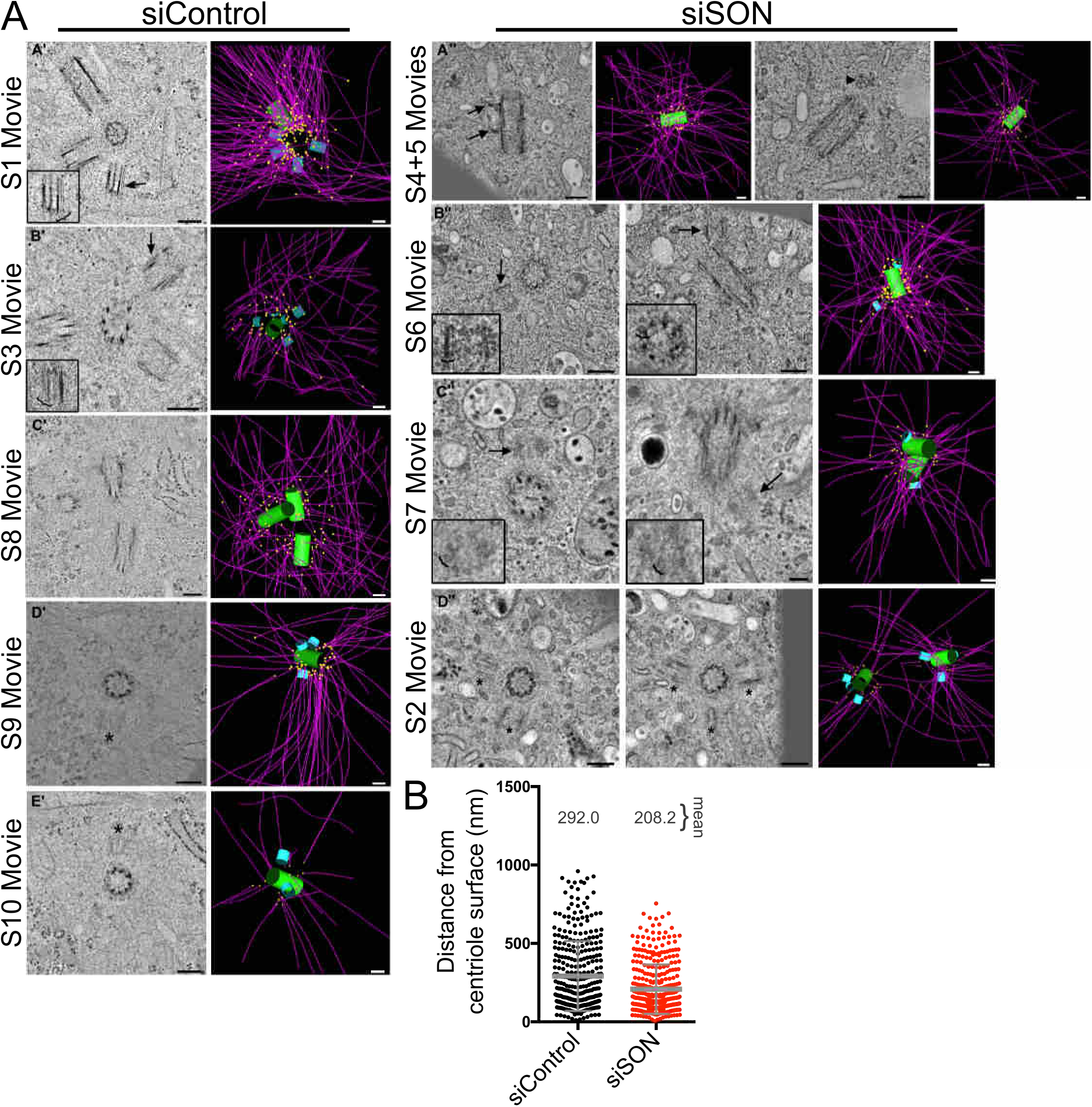
SON depletion reduces the distance of microtubule minus ends from the centriole. A) Tomographic slices and images of 3D tomographic models from all of the EM tomograms collected for this study. Images from the tomograms shown here in (A’) and (D’’) are also shown in Figure 3. Images from the tomograms in (B’) and (A’’) are shown in Figure 7. 3D models show mature centrioles in green, procentrioles in blue, microtubules in magenta and microtubule minus ends in yellow. Corresponding supplemental movies are indicated. siControl tomograms: (A’) the arrow indicates the procentriole whose microtubule triplet blades are displayed in a longitudinal view in the inset (bracket). The 3D model view does not display one of the mature centrioles to more clearly show microtubule minus ends. (B’) The arrow indicates the procentriole whose microtubule triplet blades are displayed in a longitudinal view in the inset (bracket). (C’) Three centrioles that have fully elongated. (D’) One centrosome from an siControl cell. The asterisk indicates one of the procentrioles present in the tomographic volume. (E’) The second centromsome from the same siControl cell as shown in (D’). The asterisk indicates one of the procentrioles present in the tomographic volume. siSON tomograms: (A’’) The arrows indicate the distal and subdistal appendages of the mother centriole. The arrowhead points to an amorphous density without microtubules that may indicate an early procentriole intermediate. (B’’) In the first panel, the arrow indicates the procentriole shown in longitudinal view in the inset to point out the doublet microtubules (bracket). In the second panel, the arrow indicates the procentriole shown in the inset that was rotated to display the microtubule doublets in cross section (bracket). (C’’) The arrows indicate the procentrioles shown in the insets that were rotated to display the microtubule doublets in cross section (brackets). (D’’) The first two panels show the two mature centrioles in cross section with procentrioles indicated by asterisks. Scale bars = 200 nm. B) Distances of the microtubule minus ends to the centriolar surface. N = siControl, 336; siSON, 372 from four and five tomographic models, respectively.

**Table S1. DESeq2 results.** An Excel file displaying differential expression for genes from cells treated with siSON as compared to siControl treated cells. The second tab shows the 22 CC GO terms from this dataset when the genes from the centrosome portion of the CCDB and genes from the centriolar satellite proximity screen are included as their own terms as determined by p values < 0.05.

**Table S2. MAJIQ results.** An Excel file displaying alternative splicing for genes from cells treated with siSON as determined by the MAJIQ software package. The second tab shows the 26 CC GO terms from this dataset when the genes from the centrosome portion of the CCDB and genes from the centriolar satellite proximity screen are included as their own terms as determined by p values < 0.05.

**Table S3. rMATS results.** An Excel file displaying alternative splicing for genes from cells treated with siSON as determined by the rMATS software package. Tabs display alternative 5’ splice sites, alternative 3’ splice sites, skipped exons, retained introns and mutually exclusive exons. The last tab shows the 30 CC GO terms from this dataset when the genes from the centrosome portion of the CCDB and genes from the centriolar satellite proximity screen are included as their own terms as determined by p values < 0.05.

**Table S4. The intersection between centrosomal components or satellite components and genes alternatively spliced when SON is depleted.** Genes from the centrosome portion of the CCDB (tab 1) and genes from the centriolar satellite proximity screen (tab 2) were cross referenced with genes requiring SON for proper splicing and ordered based on differential expression. Differential expression was determined by the DESeq2 software package, and results for alternative splicing as determined by the MAJIQ software package or the rMATS software package are included.

**Movie S1. Serial tomographic slices through a volume containing an siControl rosette.** The tomographic volume as built from three, serial 250 thick sections. This centrosome is displayed in Figure 3B and Figure Supplement 7A (A’). The centrosome is close to the nucleus and numerous nuclear pore complexes can be observed in the serial tomographic slices. The model shows two mother centrioles (green), and four procentrioles (blue). Microtubules (magenta) and microtubule minus ends (yellow) are modelled. Bar = 200 nm.

**Movie S2. Serial tomographic slices through a volume containing the centrosome of an siSON cell.** A tomogram from an siSON cell shown in Figure 3B and Figure Supplement 7A (D’’). The tomographic volume as built from four, serial 250 thick sections. Vesicles are present in this region around the two mother centrioles (green) and six procentrioles (blue) present in the volume. Microtubules (magenta) and microtubule minus ends (yellow) are modelled. Bar = 200 nm.

**Movie S3. Serial, tomographic slices through a volume containing the centrosome of an siControl cell.** A partial reconstruction through a volume containing the centrosome of the siControl cell shown in Figure 7E and Figure Supplement 7A (B’). This volume was built from one, 250nm thick section and therefore the mother centriole (green) and five procentrioles (blue) are not complete in the volume. Microtubules (magenta) and microtubule minus ends (yellow) are modelled. Bar = 200 nm.

**Movie S4. Serial tomographic slices through a volume containing one centrosome of an siSON cell.** This centrosome is shown in Figure 7E and Figure Supplement 7A (A’’ left). The tomographic volume as built from two, serial 250 thick sections. The model shows 1 mature centriole (green) with distal and subdistal appendages present. Microtubules can be seen originating at appendages (modelled in magenta with minus ends shown in yellow). Bar = 200 nm.

**Movie S5. Serial tomographic slices through a volume containing one centrosome of an siSON cell.** This is the second centrosome from the same cell depicted in Movie S4 and is shown in Figure Supplement 7A (A’’ right). The tomographic volume as built from two, serial 250 thick sections. No detectable procentrioles were observed, although a mass of electron dense material was observed adjacent to one of the mother centrioles. The centriole is modelled in green, the microtubules in magenta and the microtubule minus ends in yellow. Bar = 200 nm.

**Movie S6. Serial tomographic slices through a volume containing two centrosomes of an siSON cell.** The tomographic volume as built from two, serial 250 thick sections and shown in Figure Supplement 7A (B’’). Two mature centrioles (green) are observed, one of which has distal and subdistal appendages. Vesicles are present throughout the volume. The volume does not contain the entire length of one of these centrioles. Each mature centriole has a procentriole associated with it (blue). Microtubules (magenta) and microtubule minus ends (yellow) are also modelled. Bar = 200 nm.

**Movie S7. Serial tomographic slices through a volume containing two centrosomes of an siSON cell.** The tomographic volume as built from four, serial 250 thick sections shown in Figure Supplement 7A (C’’). Two mature centrioles (green) are observed, one of which has distal and subdistal appendages. Vesicles are present throughout the volume. Each mature centriole has a procentriole associated with it (blue). Microtubules (magenta) and microtubule minus ends (yellow) are also modelled. Bar = 200 nm.

**Movie S8. Serial tomographic slices through a volume containing three elongated centrioles in an siControl cell.** The tomographic volume as built from four, serial 250 thick sections and displayed in Figure Supplement 7A (C’). Three fully elongated centrioles are observed (green). Microtubules (magenta) and microtubule minus ends (yellow) are also modelled. Bar = 200 nm.

**Movie S9. Serial tomographic slices through a volume containing one of two centrosomes in an siControl cell.** The tomographic volume as built from three, serial 250 thick sections and displayed in Figure Supplement 7A (D’). The second centrosome can be seen in Movie S10. One mature centriole (green) and three procentrioles (blue) are present in the volume. Microtubules (magenta) and microtubule minus ends (yellow) are also modelled. Bar = 200 nm.

**Movie S10. Serial tomographic slices through a volume containing the second of two centrosomes in an siControl cell.** The tomographic volume as built from two, serial 250 thick sections and displayed in Figure Supplement 7A (E’). One mature centriole (green) and two procentrioles (blue) are present in the volume. The other centrosome from this cell is shown in Movie S9. Microtubules (magenta) and microtubule minus ends (yellow) are also modelled. Bar = 200 nm.

